# Characterizing shared and distinctive molecular phenotypes across motor regions in ALS with and without TDP-43 pathology in a veteran cohort

**DOI:** 10.64898/2026.08.28.747944

**Authors:** Patricia Hayes Doyle, Shiva Kazempour Dehkordi, Timothy C. Orr, Xuehan Sun, Megan S. Pater, Frederick J. Arnold, Cindy V. Ly, Miranda E. Orr

**Author notes:** Authors contributed equally (Patricia Hayes Doyle, Shiva Kazempour Dehkordi). Corresponding Author: Miranda E. Orr, 4370 Duncan Avenue, St. Louis, MO 63110.

## Abstract

Amyotrophic lateral sclerosis (ALS) is a fatal neurodegenerative disease characterized by progressive dysfunction and loss of upper and lower motor neurons. Although motor neuron degeneration ultimately drives paralysis, neuronal dysfunction may precede cell death by a prolonged interval, suggesting that vulnerable neurons engage stress-adaptive programs that permit survival despite impaired function. Cellular senescence represents one such persistent stress response and has increasingly been implicated in neurodegenerative disease, including disorders associated with TDP-43 pathology. Here, we investigated whether senescence-associated molecular states are present in vulnerable motor neurons in ALS and whether they differ according to anatomical region and phosphorylated TDP-43 (pTDP-43) pathology. Postmortem primary motor cortex, cervical spinal cord, and lumbar spinal cord were obtained from the Department of Veterans Affairs Biorepository Brain Bank from individuals with ALS classified as pTDP-43-positive or pTDP-43-negative, together with non-ALS controls. Targeted bulk transcriptomic profiling was combined with GeoMx Digital Spatial Profiling of individual motor neurons to characterize disease-, region-, and pathology-associated molecular phenotypes while preserving anatomical context. Across ALS cases, we identified alterations in pathways related to cell-cycle regulation, RNA processing, mitochondrial function, proteostasis, inflammation, and synaptic signaling. These signatures varied by anatomical region and pTDP-43 status, indicating substantial heterogeneity in the molecular response to ALS pathology. Despite these differences, both ALS groups exhibited convergent proteomic and transcriptomic features associated with cellular senescence. These findings identify senescence-associated molecular states within vulnerable neuronal populations in ALS and support a model in which persistent stress adaptation may permit neuronal survival while contributing to progressive cellular dysfunction. This spatially resolved analysis links neuronal phenotype to anatomical and pathological context and supports further evaluation of senescence-associated pathways as therapeutic vulnerabilities in ALS.

## Introduction

Amyotrophic lateral sclerosis (ALS) is a progressive neurodegenerative disease characterized by the loss of upper (UMN) and lower motor neurons (LMN), resulting in muscle weakness, bulbar dysfunction, and ultimately, fatal respiratory failure. ALS prominently affects the corticospinal tract^1–3^. Approximately half of corticospinal projections originate in the primary motor cortex^4,5^, where large layer V pyramidal neurons, including Betz cells, are particularly vulnerable to degeneration^6,7^. Lower motor neurons in the anterior horn of the spinal cord are likewise selectively affected in ALS^8,9,10,11^. This anatomically distributed pattern of degeneration suggests that molecular responses to ALS may differ across vulnerable neuronal populations and anatomical regions of the motor system.

Twin studies estimate ALS heritability at 40-60%^12–14^, yet only ∼10% of cases are associated with a known pathogenic mutation. Most ALS therefore occurs without a defined genetic cause, implicating additional environmental and non-genetic contributors to disease susceptibility. Military service is associated with increased ALS incidence ^15–19^, although neither service-related factors nor known neurotoxic exposures fully explain this increased risk^15,17,20,2115,22–24^. The availability of well-characterized postmortem tissue from veterans with ALS therefore provides an opportunity to examine molecular mechanisms underlying largely sporadic disease.

A well-established pathological feature of ALS is cytoplasmic mislocalization of TAR DNA-binding protein 43 (TDP-43), a predominantly nuclear RNA-binding protein. Loss of normal nuclear TDP-43 function and its cytoplasmic accumulation disrupt RNA processing and, protein homeostasis, and axonal transport^25,26^. In postmortem ALS cases, phosphorylated TDP-43 (pTDP-43) pathology follows a stereotyped anatomical distribution across motor and subsequently extra-motor regions, consistent with spread through connected neural networks ^27,28^. Although TDP-43 pathology is reported in up to 97% of ALS cases^12–14^, its relationship to neuronal vulnerability is not uniform across ALS subtypes or anatomical regions ^24,2927^. Rare cases with prolonged disease duration also lack detectable TDP-43 pathology ^24^, raising the possibility that TDP-43 represents one of several molecular routes to motor-neuron dysfunction. Comparing vulnerable neurons from ALS cases with and without pTDP-43 pathology may therefore distinguish TDP-43-associated changes from molecular states shared more broadly across ALS.

Cellular senescence provides one potential explanation for the prolonged dysfunction and survival of stressed motor neurons in ALS^24,30^. Senescence is a persistent cellular stress response in which damaged cells resist apoptosis while acquiring changes in morphology, macromolecular integrity, metabolism, proteostasis, and inflammatory signaling^31^. In postmitotic neurons, senescence cannot be defined by loss of replicative capacity and instead requires convergent molecular changes and associated phenotypic evidence^32–34^. Our prior work linked tau protein aggregation to a senescence-associated neuronal state and identified *CDKN2D*/p19-expressing neurons with neuropathology and characteristic senescence-associated features in postmortem human brain^35,36^.Several processes implicated in ALS, including chronic cellular stress, mitochondrial dysfunction, inflammatory signaling, altered RNA and protein homeostasis, and survival despite profound cellular dysfunction, overlap with features of cellular senescence^37–41^. Consistent with this possibility, senescence-associated p16– and p21 have been reported in ALS/MND suggesting astrocyte senescence and neuronal cell cycle dysregulation^42^. Additionally, conditional expression of cytoplasmic TDP-43 in mice linked TDP-43 pathology to DNA repair, inflammation and neuronal senescence^30^. Together these observations support testing whether senescence-associated molecular programs occur in vulnerable human motor neurons and whether they are related to TDP-43 pathology. This question is also therapeutically relevant because reducing senescent cell burden has improved neurodegeneration phenotypes in models of tauopathy, Alzheimer’s disease and Parkinson’s disease^36,43–45^.

The objective of this project was to assess cellular senescence as a potential mechanism associated with ALS in the presence or absence of aberrant TDP43 in postmortem tissue from veterans, in the absence of disease-causing mutations. We performed targeted bulk transcriptomic profiling and spatial proteomic profiling on matched tissues to characterize molecular features of ALS tissue in the motor cortex and two levels of the spinal cord (cervical and lumbar). We captured and quantified proteomic signatures in individual motor neurons while preserving their anatomical context and characterized them according to disease group, TDP-43 subcellular localization, and senescence status. By resolving these signatures at the level of individual vulnerable neurons, we aimed to identify disease-relevant processes not detectable by bulk tissue analyses, providing insight into mechanisms of ALS currently undefined or understudied. In this cohort, we identified distinct differences in disease progression according to TDP-43 status. We also found convergent molecular signatures in both types of ALS associated with senescence, with distinctions according to TDP-43 status, tissue region, and pTDP-43 levels, underlining differences in disease-associated molecular state and coordination of cellular response to ALS.

## Methods

### Human postmortem cases, tissue processing, and neuropathologic characterization

The Department of Veterans Affairs Biorepository Brain Bank (VA BBB) prospectively collects, processes, and distributes central nervous system tissue for research on ALS and other neurological disorders under Institutional Review Board approval (IRB #1938) and VA Merit Review Award BX002466. Available clinical and demographic variables included age at death, sex, ethnicity, postmortem interval (PMI), cause of death, ALS subtype (sporadic or familial), disease duration, neuropathologic diagnosis, and TDP-43 stage, where applicable. Individuals with ALS underwent longitudinal clinical assessment using the ALS Functional Rating Scale–Revised (ALSFRS-R)^46^ at enrollment and approximately every six months thereafter.

Brain and spinal cord tissues were collected, fixed, processed, and neuropathologically characterized according to the standardized VA BBB protocol as previously described^47^. Briefly, one cerebral hemisphere was fixed while the contralateral hemisphere was dissected and snap-frozen. Following fixation, representative brain and spinal cord regions were processed into FFPE tissue blocks for histopathologic evaluation. Baseline neuropathologic assessment included phosphorylated TDP-43, phosphorylated tau, α-synuclein, ubiquitin, and amyloid-β pathology, and ALS neuropathologic staging was assigned according to established criteria^24,28,48^.

Postmortem central nervous system tissue and associated clinical metadata were obtained from the VA BBB. Cases were selected a priori to include 12 TDP-43-positive ALS, 12 TDP-43-negative ALS, and 12 non-ALS control donors (total n = 36, **Fig. 1**). TDP-43 positive cases (ALS-TDP-43(+)) were defined as having cytoplasmic pTDP-43 found at any level of the CNS. ALS and control groups were matched by age and sex, and cases with known pathogenic ALS-associated mutations were excluded.

**Figure 1.**
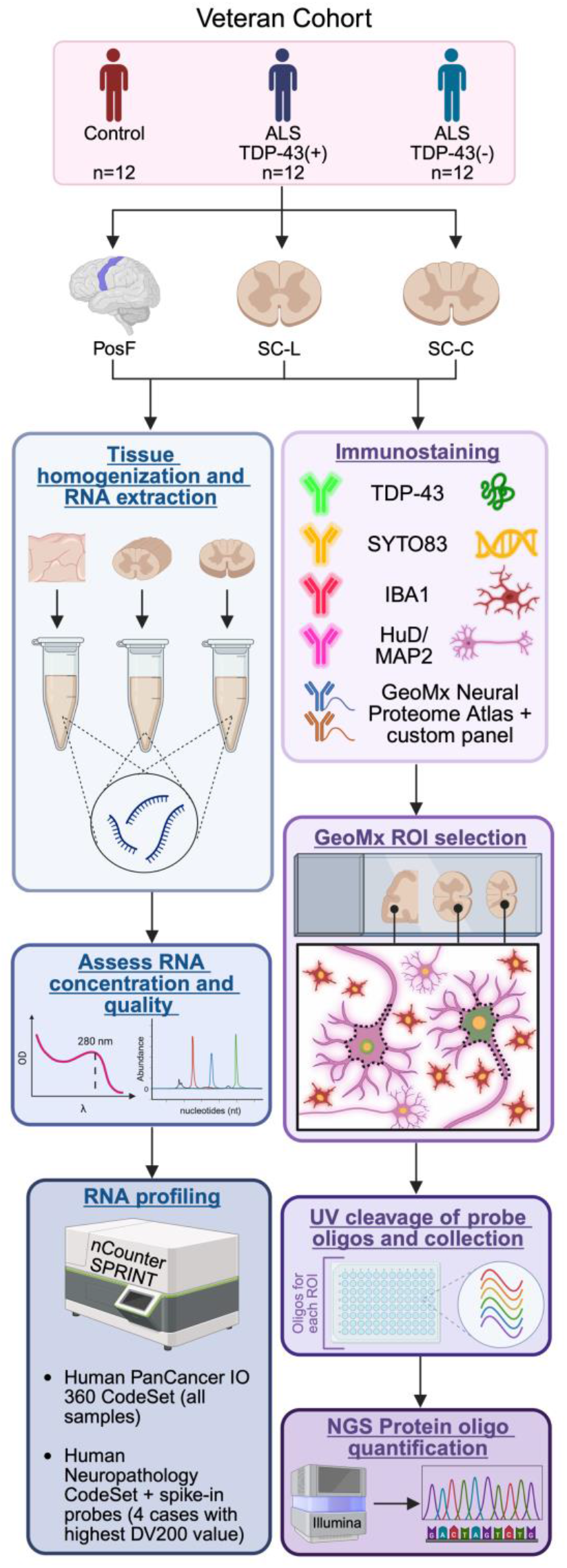
Study design and experimental workflow for transcriptomic and spatial proteomic profiling of ALS spinal cord tissue. Veteran tissue from the posterior frontal cortex (PosF, containing primary motor cortex), lumbar spinal cord (SC-L), and cervical spinal cord (SC-C) was obtained from the VA Biorepository Brain Bank (*top*). The RNA profiling workflow (*left*) utilized targeted panels to assess inflammatory and neuropathology-related transcriptomic changes across each tissue. The spatial proteomic workflow (*right*) used TDP-43, nucleic acid (SYTO 83), microglial (IBA1) and neuronal (HuD, MAP2) morphology markers and UV-cleavable protein probes to profile motor neurons of interest for protein indicators of senescence. Next-generation sequencing of oligonucleotides enabled quantification of proteins of interest.

For each donor, tissue was obtained from three anatomically matched regions: the primary motor cortex from the posterior frontal cortex (Brodmann area 4, PosF), cervical spinal cord (SC-C), and lumbar spinal cord (SC-L). Formalin-fixed paraffin-embedded (FFPE) tissue blocks from each region were used for spatial proteomic profiling, while matched frozen tissue from the same donors and anatomical regions was used for targeted RNA expression profiling using the NanoString nCounter platform. In total, the study analyzed 108 FFPE specimens and 108 matched frozen specimens across the three anatomical regions.

### RNA isolation and nCounter analysis

Total RNA was isolated from ≥20 mg of frozen tissue from the PosF, SC-C, and SC-L using the RNeasy Midi Kit according to the manufacturer’s protocol (Version: Apr. 2021; **Fig. 1**, left). Tissue samples were homogenized in Buffer RLT supplemented with 1% 2-mercaptoethanol using pellet pestles, followed by additional homogenization by passage through 21-gauge needles. RNA concentration was determined using a NanoDrop 8000 spectrophotometer, and RNA quality was assessed by DV200 using an Agilent TapeStation 4200. All reagent and equipment information is provided in **Tables S1** and **S2.**

RNA abundance was quantified using the nCounter Human PanCancer IO 360 CodeSet (on an nCounter SPRINT Profiler according to the manufacturer’s recommendations). RNA input for each reaction was adjusted based on the measured DV200 value to account for differences in RNA integrity. For each anatomical region, the four specimen with the highest DV200 values per group were selected for additional profiling using the nCounter Human Neuropathology CodeSet supplemented with ten custom spike-in probes targeting *CDKN2D, MAPT* (exon 4a), *ANK3, SPTN4, MAP6, CHMP7, POM121, KPNA1, SRSF6,* and *HNRNPM*.

### GeoMx Slide Preparation

FFPE tissue sections were prepared for GeoMx Digital Spatial Profiler (DSP) protein profiling according to the manufacturer’s protocol (Version: MAN-10150-07, Sept. 2025; **Figure 1**, right) and our previously published methods^49–51^. Briefly, slides were deparaffinized, subjected to heat-induced antigen retrieval, and processed using the GeoMx DSP Protein Assay workflow. Minor modifications to the standard protocol were implemented to facilitate removal of excess paraffin present on tissue sections received from the VA Biorepository Brain Bank and to improve morphology visualization. Specifically, slides were baked for 45 minutes at 60°C before deparaffinization, and an additional CitriSolv wash was incorporated following a brief immersion in 100% ethanol. We also found that preheating Citrate Buffer prior to antigen retrieval improved cellular morphology. All subsequent steps followed the manufacturer’s recommended protocol. Reagents, consumables, and equipment are listed in **Tables S1** and **S2**.

Protein probes included the GeoMx Core, Neural Proteome Atlas, and a custom panel of 21 proteins (detailed in **Table S3**). Fluorescent antibodies for morphological and histological markers included IBA1 (1:100, activated microglia), HUD (1:100) and MAP2 (1:100, neurons), SYTO83 (1:15000, nucleic acid), and TDP-43 (ALS-associated inclusions). The same TDP-43 antibody was used for all slides. For 48 of the 108 slides, the antibody was conjugated using the Flexlinker kit and diluted 1:10 to enhance signal. The remaining slides were stained using the TDP-43 antibody without additional Flexlinker conjugation (1:100).

To minimize batch effects, slides were processed in 10 groups; 9 included slides from all neurological regions and diagnosis statuses. Slides were added to the GeoMx Digital Spatial Profiler in groups of 3, with well A1 of the control slide being reserved for non-template control and 32 AOIs assigned per remaining slide.

### Motor Neuron Identification, AOI Selection, and Single-Cell Mask Generation

Motor neurons were identified by HuD and MAP2 immunofluorescence together with their characteristic morphology. A minimum of 31 motor neurons was selected from each tissue section for each pathological group (control, TDP-43-negative ALS, and TDP-43-positive ALS). When fewer than the target number of morphologically identifiable motor neurons were present, all available motor neurons were analyzed. Exact sample sizes for each analysis are provided in the corresponding figure legends.

Following whole-slide imaging on the GeoMx Digital Spatial Profiler (DSP), areas of interest (AOIs) were manually drawn around individual motor neurons using fluorescence morphology markers. Each AOI was intentionally drawn larger than the target neuron to ensure complete cellular capture and to provide sufficient surrounding area for subsequent image-based segmentation. AOI images were exported from the GeoMx DSP software and processed in ImageJ (FIJI)^52^ using a custom macro (MaskMacro) developed to generate binary segmentation masks for individual motor neurons. The workflow used automated thresholding and particle-based segmentation to identify neuronal cell bodies, as shown below:

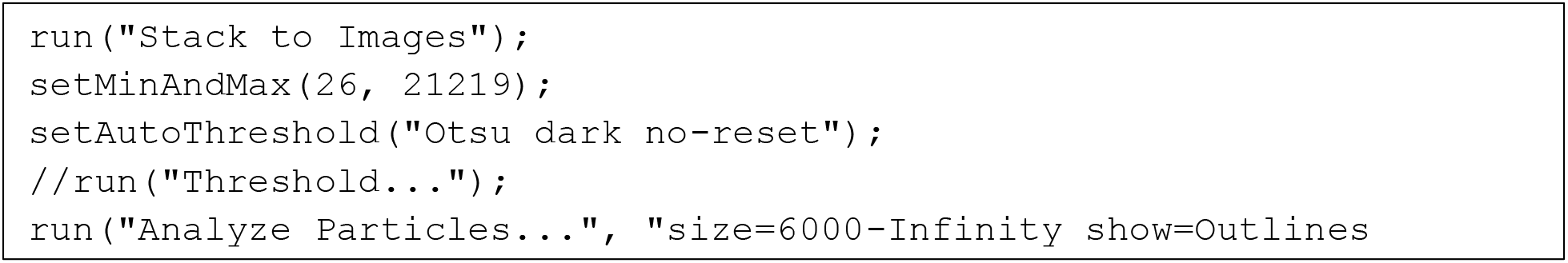

Minor adjustments were made to threshold and particle-size parameters when necessary to accommodate differences in staining intensity, neuronal size, or image quality. Segmentation masks were visually reviewed for accuracy before being imported back into the GeoMx DSP software. The imported binary masks were used as custom areas of illumination, restricting UV photocleavage to the segmented motor neuron while excluding surrounding neuropil and neighboring cells. This single-cell masking workflow was applied to approximately **3,500 individual motor neurons** across the PosF, SC-C, and SC-L. Representative images are reflected in **Fig. 2**.

**Figure 2.**
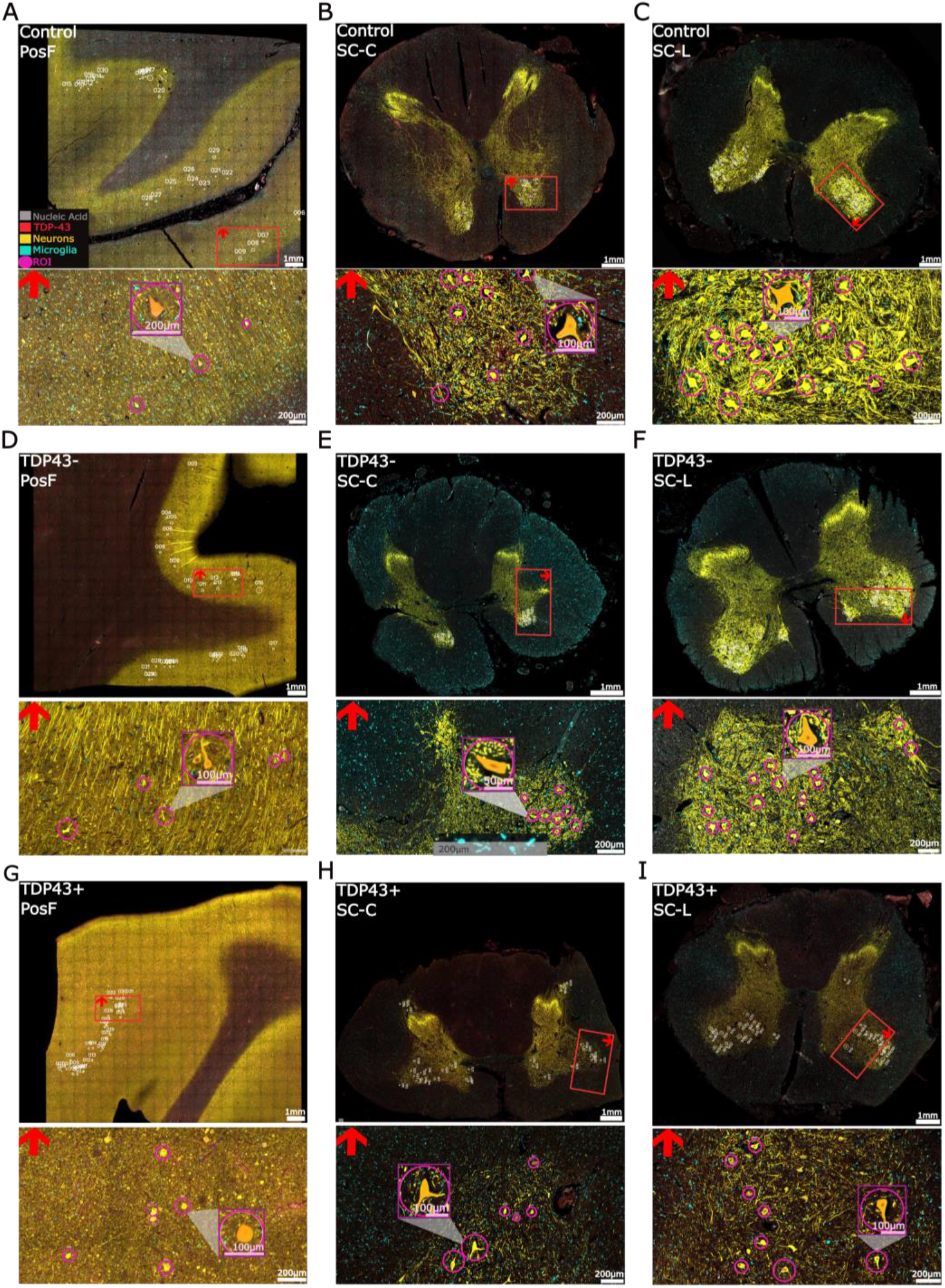
Representative GeoMx images for morphology visualization and AOI selection. (**A-I**). Representative GeoMx images from three disease condition groups: Control (**A-C**), TDP-43-negative (**D-F**), or TDP-43-positive (**G-I**). Each row of images corresponds to a single representative case, with images from the same case shown across three columns. Columns correspond to the Posterior Frontal Cortex (PosF, motor cortex, left), cervical spinal cord (SC-C, middle), and lumbar spinal cord (SC-L, right). For each region, a broad-field image is shown above the corresponding zoomed-in view, with the red box indicating the area shown in the zoomed image. Spinal cord images were oriented with the dorsal horn toward the bottom of the broad image, with a red arrow in the upper left of the zoomed image indicating orientation. Regions of interest (AOIs) were circled in pink and the following morphology markers were used to select AOIs, pseudocolored as indicated (**A, top**): nucleic acids (grey), TDP-43 (red), neurons (yellow), and microglia (cyan). AOI masking is demonstrated with an orange overlay for one neuron per image, as indicated in the purple zoom square.

### NGS Protein quantification

Photocleaved oligonucleotide tags collected from the GeoMx Digital Spatial Profiler (DSP) were prepared for next-generation sequencing according to the manufacturer’s protocol (GeoMx DSP NGS Readout, MAN-10153-01) with minor modifications. To minimize technical variability, libraries were prepared in batches of 2–4 DSP collection plates. Oligonucleotides were dried and rehydrated in 30μL, rather than the recommended 80μL, due to the small AOI size. Sequencing libraries were generated using the GeoMx NGS Master Mix and ProCode primers according to the manufacturer’s recommendations, aside from using 4μL (rather than the recommended 2μL) of DSP aspirate to maximize protein detection in our small surface area. PCR products were purified using AMPure beads, quantified using the Qubit dsDNA High Sensitivity Assay, and normalized before sequencing with 1.5% PhiX spike-in.

Libraries with compatible ProCodes were pooled and sequenced on an Illumina NextSeq 1000/2000 platform using P2 or P4 flow cells according to the manufacturer’s recommendations. Complete reagent and equipment information is provided in **Tables S1** and **S2**, and thermocycler conditions are listed in **Table S4**.

### Demographic Statistics

Initial demographic analyses were performed in R (v4.6.1)^53^. Data visualization was performed using ggplot2 (v4.0.3)^54^, ggsignif (v0.6.4)^55^, and showtext (v0.9-8)^56^, and data manipulation was performed using dplyr (v1.2.1)^57^ and tidyr (v1.3.2)^58^.

Kruskal-Wallis tests (`kruskal.test`)^59^ were performed to assess significant differences in characteristics measured across all three groups (age at death, PMI, RNA Integrity Number [RIN], DV200, pH). Wilcoxon rank-sum tests (`wilcox.test`)^60,61^ were performed to assess significance between groups in disease-specific characteristics (age at diagnosis, disease duration). Significance was defined at α=0.05.

Anterior horn cellular degeneration was classified as an ordered categorical variable with four levels: mild, moderate, marked, and severe loss. The distribution of severity of anterior horn cell loss between ALS-TDP-43(+) and TDP-43-negative was evaluated using Wilcoxon-rank-sum test (`wilcox.test`)^60,61^. To assess whether ALS-TDP-43(+) cases had a greater likelihood of exhibiting greater severity of cell loss in the anterior horn than ALS-TDP-43(-) cases, a proportional-odds logistic regression was fitted using the `polr` function from the MASS package (v7.3.66)^62,63^. Severity was specified as an ordinal categorical outcome, with TDP-43 status being assigned as the predictive variable and the TDP-43-positive group specified as the reference group. The analysis followed the approach described by McNulty^62^.

To assess variation in RNA quality prior to transcriptomic analyses, RIN and DV200 measures were analyzed across our acquired tissues (PosF, SC-C, SC-L) and compared to reported values from occipital tissue. Cases with missing data across tissue regions were excluded for these analyses. Measurements from the same individual were treated as repeated observations. Aligned rank transform (ART) mixed-effects models were fitted using the ARTool (v0.11.2)^64^ and lme4 (v2.0.6)^55^ packages. Disease group, region, and the group x tissue region interaction were treated as fixed independent variables and case as a random intercept. When significant effects were identified, post-hoc comparisons using ART-C (aligned rank transform contrast, `art.con`)^65^ were performed, using the Holm^66^ approach to correct for multiple comparisons. Regional pairwise comparisons within each disease group were performed for significant disease group x tissue region interactions using estimated marginal means (emmeans, v2.0.4)^67^ from the ART interaction model (`artlm`), with Holm^66^ adjustment for multiple comparisons.

### NCounter Analyses

#### NCounter data processing and normalization

We profiled bulk tissue on the NanoString nCounter platform^68^ using two panels: a 628-gene Cancer IO and immune panel and a Neuropath panel, each carrying NanoString’s standard positive (POS_A–F), negative (NEG_1–8), and housekeeping control probes.

We ran QC and normalization in R (v4.5.1)^53^ with custom functions and followed Nanostring’s Gene Expression Analysis Guidelines separately for each panel and within two tissue groupings: posterior frontal cortex (PosF) and spinal cord (cervical, “SC-C,” and lumbar, “SC-L,” pooled as “SpinalCord”).

Positive-control linearity QC: For each sample, we regressed log2 counts of the five highest-concentration positive control probes (POS_A–E; POS_F excluded as below the assay’s limit of detection) against log2 known concentration and flagged samples with R² < 0.95.

<u>Negative-control background</u>: For each sample, we checked each of the 8 negative control probes against the average of the other 7. A probe that came in more than 3-fold higher than that average was excluded from the sample’s background estimate. We defined background as the mean plus two standard deviations of the surviving negative-control probes.

Limit-of-detection (LOD) QC: We flagged a sample with a failing LOD if its POS_E count (the lowest positive-control concentration, 0.5 fM) did not exceed its cleaned background.

<u>Background thresholding</u>: We dropped the negative control probes and raised any remaining probe count below a sample’s background up to that background level. Thresholding was favored in this dataset than subtraction as we would not inflate fold-change estimates for genes being near background.

<u>Positive-control normalization</u>: We computed a per-sample scale factor from the geometric mean of raw POS_A-E count as grand mean of all samples’ geometric means, divided by each sample’s own. We then applied it to the background-thresholded counts. We flagged samples with a factor outside of range [0.3, 3].

<u>Housekeeping normalization</u>: We first screened candidate housekeeping genes for stability using the pairwise variation algorithm^69^. In this method, we iteratively dropped the least stable candidate until every remaining gene’s M-value was ≤ 1.5. We excluded *GUSB* and *FAM104A* from the candidate pool, since both correlated poorly with the other housekeeping genes. We then computed a per-sample scale factor from the geometric mean of the retained stable genes and applied it on top of the positive-control-normalized counts. Samples with a factor outside [0.1, 10.0] were flagged.

#### Multifactor differential expression analysis

We tested differential expression between disease condition groups (Control, ALS-TDP-43(-), ALS-TDP-43(+)) separately within each panel and tissue grouping, using DESeq2 package (v1.50.2)^70^. We fit the negative-binomial model on raw counts using `DESeqDataSetFromMatrix(), DESeq()` functions. Instead of DESeq2’s default median-of-ratios size factors, we set `sizeFactors()` to use this cohort’s own combined positive-control times housekeeping scale factor. We inverted this value since DESeq2 divides by size factor, but our normalization above multiplies by scale factor.

We fit an intercept-based additive design, `Disease Condition + Tissue` and dropped `Tissue` for PosF since it is constant. We called a gene differentially expressed per coefficient at Benjamini-Hochberg adjusted p < 0.05 and absolute value of log2 fold-change > 0. we also fit `Tissue + Disease Condition + Tissue:Disease Condition` to ask whether the Disease Condition effect differs between cervical (SC-C) and lumbar (SC-L) cord, testing the interaction coefficient(s) jointly with a BH-corrected LRT against a reduced model that kept both main effects. For each comparison we used the ComplexHeatmap package ^71^to visualize the heatmaps.

#### Pathway enrichment analysis

For every DE contrast (individual coefficient or combined LRT result) with more than 5 differentially expressed genes, we ran gene set enrichment analysis (GSEA)^72^ using clusterProfiler (v4.18.4)^73^ in R. We ranked the full detected gene list by signed −log10(p-value), with sign taken from the log2 fold-change direction. We kept terms at Benjamini-Hochberg adjusted p < 0.05 and produced per-database dotplots, ridgeplots, NES bar plots, and enrichment maps with enrichplot (v1.30.5)^73^ and ggplot2 (v4.0.3)^54^. To summarize a contrast’s signal across databases in one figure, we built combined bubble plots (custom ggplot2 script) that pool the top N significant GSEA terms per database (adjusted p < 0.05; N = 7–10 depending on the figure) onto a single shared axis.

### GeoMx protein analysis

#### GeoMx protein QC and normalization

We ran QC and normalization steps in R (v4.5.1)^53^ with custom functions. We excluded AOIs from a confirmed bad case (n=1) and bad sequencing runs (n=2).

Normalization was done in two steps: First, we used four stable IgG isotype contrl probes (Hmr IgG, Rb IgG, Ms IgG1, Rt IgG2a) and checked their stability across AOIs, their concordance with each other, and correlation AOI area and dropped any that failed. We next computed each AOIs IgG geometric mean (geomean) and flagged AOIs with a log10(IgG geomean) Z-score more than 3 as high-background. We next used an RLE-based outlier check in a way that an AOI’s median log-ratio to the panel-wide median across signal proteins should not exceed 3 in absolute value. Finally, we applied the third check using a low-signal threshold. We computed an AOI’s total signal and removed any with that value more than 2 standard deviations (SD) below the mean. We then set a per-AOI limit of quantification (LOQ) from each AOI’s IgG geomean and geoSD, and kept proteins if their raw count passed that LOQ in at least 1% of AOIs. On the proteins that passed, we scaled each AOI by the ratio of the global IgG geometric mean to that AOI’s own. We call this normalization NegNorm. This approach, and the specific QC checks above, follow published GeoMx normalization practices^74^ for protein data.

**Table 1.** AOIs, cases, and proteins retained per tissue after normalization.

| Tissue | AOIs (cases) | AOI QC-flagged | Proteins retained (%) |
| --- | --- | --- | --- |
| PosF | 1,009 (32) | 36 (3.6%) | 649 (96%) |
| SC-C | 973 (31) | 96 (9.9%) | 637 (94%) |
| SC-L | 952 (31) | 36 (3.8%) | 636 (94%) |

Second, this cohort also carries a shared per-AOI loading confound meaning that every protein pairwise-correlates at r = 0.4–0.95 on `log2NegNorm` values. Before fitting any statistical models on the `log2NegNorm,` we subtracted each AOI’s own mean across the full detected panel, producing a per-AOI ratio matrix. When a model’s outcome was itself one of the marker proteins, we left that marker out of the averaging step, so it wasn’t used to correct itself (e.g., pTDP-43, TDP-43).

#### Differential expression and pathways enrichment analyses

We tested differential protein expression between Disease Condition levels (Control, ALS-TDP-43(-), ALS-TDP-43(+)) separately per tissue (PosF, SC-C, SC-L) and a pooled spinal cord with Tissue added as a fixed-effect covariate. We fit a mixed linear model with limma (v3.66.0)^75^, using the `duplicateCorrelation()`^76^ function to estimate and block on the intra-case correlation induced by multiple AOIs per case. Then `eBayes(tred=TRUE)` was used to obtain moderated statistics for every pairwise contrast. We called a protein significant at BH adjusted p-value less than 0.05 per contrast.

For every contrast, we ran gene set enrichment analysis (GSEA, as described under nCounter data analysis section). We mapped each protein name to its HUGO Gene Nomenclature Committee (HGNC) Symbol^77^ provided by Bruker Spatial Biology. We used ReactomePA (v1.54.0)^78^ and ranked the full protein list by signed −log10(p-value). Gene sets were restricted to size 10–500 and terms retained at Benjamini-Hochberg adjusted p-value < 0.05.

#### Eigenprotein analysis

To summarize the panel’s ∼630 proteins into a smaller number of biologically interpretable axes, we scored each AOI on a set of curated biology-informed protein modules (**Table 2**). An eigenprotein for a given set was set as the first principal component (PCA). For each tissue and each module with at least 3 detected members, we calculated the eigenprotein and flipped the PC1’s sign as needed, so a higher score means higher average module expression.

**Table 2.** Curated protein modules for eigenprotein analysis.

| Program | n | Members |
| --- | --- | --- |
| Senescence | 11 | p21; 53BP1; p18 INK4c/CDKN2C; p19 INK4d_CDKN2D; E2F4 (not detected in SC-C); ASK1; JNK2; MAP4K4/NIK; p53 (phospho S392) EP155Y; p53 (phospho S6); p53 E26 |
| Metabolism | 11 | Glucose Transporter GLUT4; Glucose Transporter GLUT3 + GLUT14; VDAC2; VDAC1/Porin; PKM; SLC1A5/ASCT2; GLUD1 + GLUD2; Cytochrome C; Nrf2 (phospho S40); SDHB; OPA1 |
| OXPHOS | 20 | NDUFA10; NDUFAB1; NDUFB3; NDUFB8; NDUFB9; NDUFB10; NDUFB11; NDUFS3; NDUFS6; NDUFS8; DAP13/NDUFA12; SDHB; UQCRB; UQCRC2; UQCRH; COX IV; COX5A; COX5B; COX6B1; COX7B |
| Cell cycle | 7 | Cdk1-2-3-5; Cyclin A1; Cyclin B2/CCNB2; Cyclin D3/CCND3; CDK5RAP3; p21; p18 INK4c/CDKN2C |
| Autophagy | 8 | ATG3; AMPK beta 1; eEF1A1/EF-Tu; PARK7/DJ1; Hsc70; Retinoic Acid Receptor alpha; Huntingtin; Nrf2 (phospho S40) |
| RBPs | 11 | TDP43; hnRNP H; Matrin 3; DDX5; DDX17; RNA Helicase A; nmt55/p54nrb; RALY; ILF3; TLS/FUS; TAF15 |
| Alternative-splicing | 8 | Nova1; A2BP1/Fox1/RBFOX1; HuD; SC35; SF3A1; SF3B3; PRPF8/Prp8; TRA2B/SFRS10 |

We tested each module individually for a Disease Condition effect with a linear mixed model (`Score ∼ Disease Condition + (1|Case)`). We used lme4 (v2.0.1)^79^ and lmerTest (v3.2.1)^80^ to get a Satterthwaite F-test^81^ for the Disease Condition term and emeans (v2.0.3)^67^ for pairwise contrasts. We also asked whether the modules shift jointly by Disease Condition. For that, we averaged each module’s AOI-level score to one value per case and fit a classic MANOVA (Pillai’s trace, base R ‘stats::manova()’) on the case-level score matrix. We next followed that up with a per-program univariate ANOVA to show which module(s) drove a significant joint effect.

#### Within-case pTDP43 and TDP43 slope analysis

To ask whether increasing neuronal pTDP-43 (or TDP43) burden associates with different molecular changes depending on Disease Condition and whether that relationship differs by tissue, we fit tissue-stratified mixed models (PosF, SC-C, and SC-L each fit separately). For each tissue, we decomposed each AOI’s pTDP-43 and TDP43 signal into a within-case deviation and a case-mean term. The within-case value is the AOI’s raw signal minus its case’s mean, and the case-level average tells how far that individual AOI deviates from its own case’s average. This split matters because the within-case value averages to zero within every case, by construction. That means it can’t be confounded by anything that’s constant within a case. So any relationship we find between the within-case value and another protein has to come from AOI-to-AOI variation, not from which group the case belongs to. We then divided each tissue’s within-case values by that tissue’s own standard deviation.

For every protein (excluding the two marker rows themselves), we fit pTDPwithinZ*DiseaseCondition+ TDP43withinZ*DiseaseCondition+ pTDPcaseMean + TDP43caseMean + AreaWithin + AreaCaseMean + (1|Case) with dream()^82^ from variancePartition (v1.40.2)^83^, followed by eBayes()^84^ for moderated statistics. We extracted each Disease Condition level’s slope for pTDP-43 and for TDP43 as a linear contrast (variancePartition::makeContrastsDream()) from this single shared fit. We next classified each protein’s Disease Condition-specific sign and significance pattern into disease-stage archetypes (e.g. conserved across all three groups, ALS-TDP-43(+) only, present in both disease groups but not Controls, lost specifically in ALS-TDP-43(+), or sign-switching between ALS-TDP-43(-) and ALS-TDP-43(+)) and summarized these, together with slope heatmaps and forest plots (β ± 95% CI) for a panel of proteins flagged as candidates from earlier exploration, across all three tissues.

## Results

### Cohort and tissue overview

Demographic and clinical characteristics of the study cohort are summarized in **Table 3**. The study cohort was primarily Caucasian (29/36, 80.6%) and male (33/36, 91.7%). The control group contained the only female and all but one of the non-Caucasian participants. Median age at death was 70 years, with no significant difference between groups (p = 0.161, **Fig. 3A**), however this characteristic was listed as “unknown” for two of the control cases.

**Figure 3.**
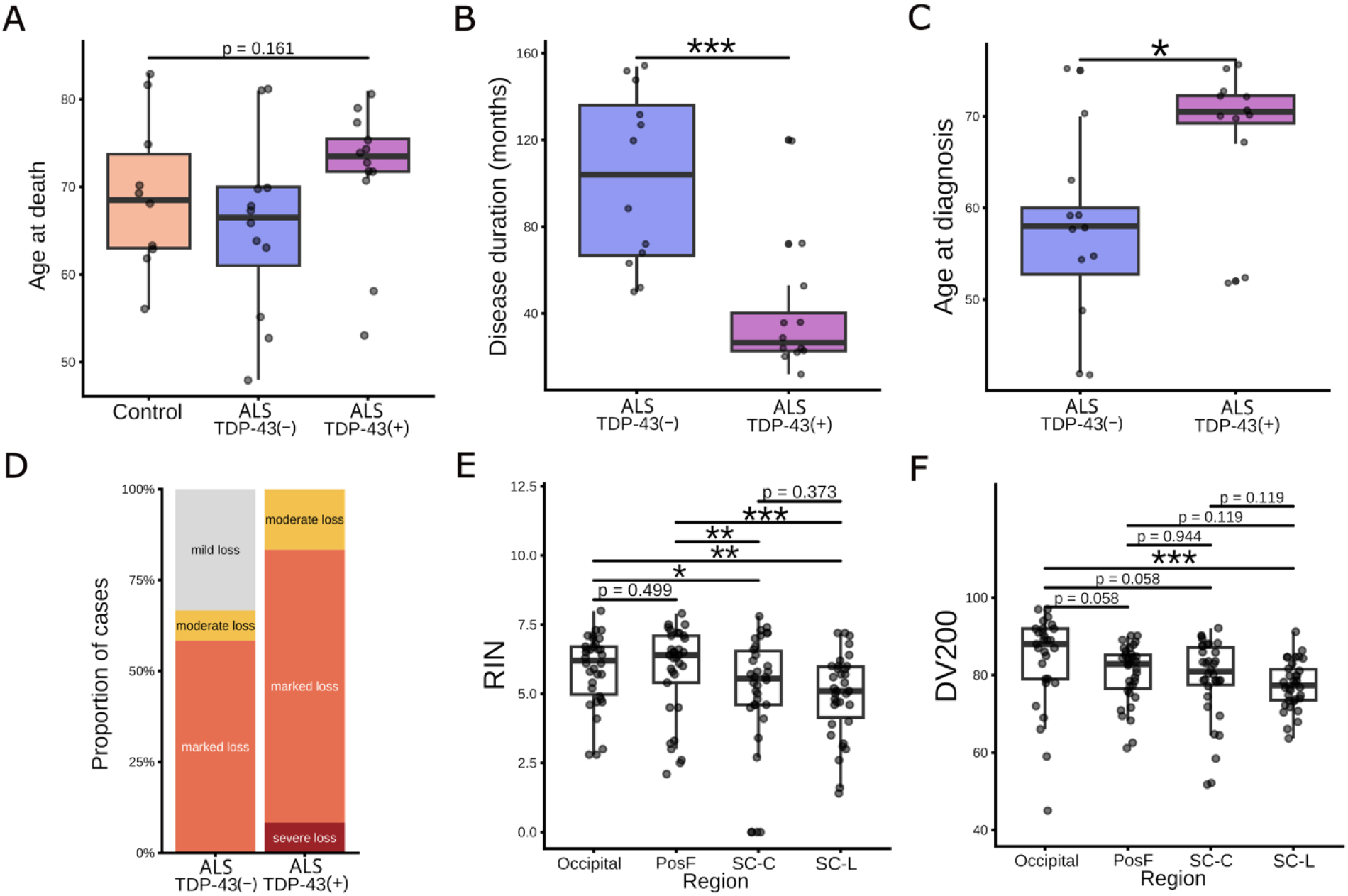
Analyses of clinical and pathological characteristics of study cohort reflect pathological grouping-specific deviations in disease progression. **(A)** Age at death across disease groups, with no significance observed (Kruskal-Wallis test, p = 0.161. **(B)** Disease duration among ALS-TDP-43(-) and ALS-TDP-43(+) ALS cases. ALS-TDP-43(-) cases had significantly longer disease duration than ALS-TDP-43(+) cases (Wilcoxon rank-sum test, ***p<0.001) **(C)** Age at diagnosis of ALS among cases with and without TDP-43. ALS-TDP-43(-) cases were diagnosed at a significantly younger age than ALS-TDP-43(+) cases (Wilcoxon rank-sum test, *p <0.05) **(D)** Distribution of anterior horn loss severity among ALS-TDP-43(+) and ALS-TDP-43(-) ALS cases. Distribution of anterior horn cell loss was not significantly different between groups (Wilcoxon-rank-sum test, p = 0.081). (**E**) RNA Integrity Number (RIN) across occipital lobe, posterior frontal cortex (PosF), and spinal cord from the cervical (SC-C) and lumbar (SC-L) levels. A significant effect of tissue region was detected (ART ANOVA, ****p < 0.0001), with Holm-adjusted post-hoc tests revealing significantly lower RIN in SC-C (*p<0.05) and SC-L (**p < 0.01) compared to occipital tissue, and significantly higher RIN in PosF compared to SC-C and SC-L (p <0.05). (**F**) DV200 percentages across tissue regions. A significant effect of tissue region (***p <0.001) was identified and Holm-corrected post-hoc tests showed significantly lower DV200 in SC-L compared to occipital tissue (****p=0.0001).

**Table 3.** Demographic, c linical, and neuropathological characteristics of study participants.

| Characteristic | Overall | Control | ALS,<br>TDP-43(-) | ALS,<br>TDP-43(+) |
| --- | --- | --- | --- | --- |
|  | *n* = 36 | *n* = 12 | *n* = 12 | *n* = 12 |
| **Sex, n (%)** |  |  |  |  |
| Male | 33 (91.7%) | 9 (75%) | 12 (100%) | 12 (100%) |
| Female | 1 (2.8%) | 1 (8.3%) | 0 | 0 |
| Unknown | 2 (5.6%) | 2 (16.7%) | 0 | 0 |
| **Ethnicity, n (%)** |  |  |  |  |
| Caucasian | 29 (80.6%) | 7 (58.3%) | 11 (91.7%) | 11 (91.7%) |
| Asian | 2 (5.6%) | 1 (8.3%) | 0 | 1 (8.3%) |
| Hispanic | 2 (5.6%) | 2 (16.7%) | 0 | 0 |
| Black/AA | 1 (2.8%) | 0 | 1 (8.3%) | 0 |
| Unknown | 2 (5.6%) | 2 (16.7%) | 0 | 0 |
| **Age at death, years** |  |  |  |  |
| Mean ± SD | 68.7 ± 9.1 | 69.1 ± 8.8 | 65.5 ± 10 | 71.6 ± 8.1 |
| Median (Q1, Q3) | 70 (63, 74.8) | 68.5 (63, 73.8) | 66.5 (61, 70) | 73.5 (71.8, 75.5) |
| Range | 48-88 | 56-88 | 48-81 | 53-81 |
| **Age at ALS diagnosis, years** |  |  |  |  |
| Mean (± SD) | 62.7 ± 10.5 | - | 57 ± 9.8 | 68.3 ± 8 |
| Median (Q1, Q3) | 65 (54.8, 71.3) | - | 58 (52.8, 60) | 70.5 (69.3, 72.3) |
| Range | 42-76 | - | 42-75 | 52-76 |
| **Disease duration, months** |  |  |  |  |
| Mean (± SD) | 70.7 ± 47.5 | - | 102.2 ± 40.6 | 39.3 ± 30.2 |
| Median (IQR) | 58 (27.8, 120) | - | 104 (66.8, 136) | 26.5 (22.8, 40.3) |
| Range | 12-154 | - | 50-154 | 12-120 |
| **Site of Onset, n (%)** |  |  |  |  |
| Upper Extremities | 9 (37.5%) | - | 4 (33.3%) | 5 (41.7%) |
| Lower Extremities | 10 (41.7%) | - | 6 (50.0%) | 4 (33.3%) |
| Bulbar | 5 (20.8%) | - | 2 (16.7%) | 3 (25.0%) |
| **ALS type, n (%)** |  |  |  |  |
| Sporadic | 17 (70.8%) | - | 5 (41.7%) | 12 (100%) |
| Familial | 3 (12.5%) | - | 3 (25%) | 0 |
| Unknown | 4 (16.7%) | - | 4 (33.3%) | 0 |
| **TDP-43 Staging** |  |  |  |  |
| Stage 2 | 1 (8.3%) | - | - | 1 (8.3%) |
| Stage 3 | 3 (25.0%) | - | - | 3 (25.0%) |
| Stage 4 | 8 (66.7%) | - | - | 8 (66.7%) |
| **Anterior Horn Cell Loss** |  |  |  |  |
| Mild | 4 (16.7%) | - | 4 (33.3%) | 0 |
| Moderate | 3 (12.5%) | - | 1 (8.3%) | 2 (16.7%) |
| Marked | 16 (66.7%) | - | 7 (58.3%) | 9 (75.0%) |
| Severe | 1 (4.2%) | - | 0 | 1 (8.3%) |

**Abbreviations:** AA, African American. **Note:** Summary statistics for continuous variables were calculated from available data. Percentages for ALS-specific variables (age at diagnosis, disease duration, ALS type) are calculated among ALS cases only (*n* = 24).

Clinical characteristics differed by TDP-43 pathology status. The site of onset was fairly similar between ALS types, with lower-limb onset being the most common presentation overall (10/24, 41.7%) and being more frequent among ALS-TDP-43(-) cases (6/12, 50%). Upper-limb onset was slightly more frequent among ALS-TDP-43(+) cases (5/12, 41.7%). The ALS TDP-43-negative group (ALS-TDP-43(-)) included both sporadic, familial, and cases with unknown ALS classification, whereas all TDP-43-positive (ALS-TDP-43(+)) cases in this cohort were sporadic and most exhibited advanced pathological staging, as described by Brettschneider *et al*.^28^ classified as stage 3+ or higher (91.7%). Notably, ALS cases negative for TDP-43 pathology exhibited a significantly younger median age at diagnosis compared with ALS-TDP-43(+) cases, by over a decade (58 vs. 70.5 years of age; p = 0.011, **Fig. 3B**) and had a significantly longer median disease duration (104 vs. 26.5 months; p = 0.0007, **Fig. 3C**).

ALS-associated degeneration predominantly affects the motor neurons within the corticospinal tract^1–3^, including the LMN in the anterior horns of the spinal cord^8,9^. Regional pathological notes of ALS cases confirmed motor system degeneration at the level of the cervical, thoracic, and lumbar spinal cord in all cases, characterized by anterior horn cell loss or degeneration of the anterior gray matter or corticospinal tracts. Prominent gliosis was also noted in several cases. To compare disease staging between ALS cases with and without TDP-43 pathology, we used overall anterior horn cell loss severity as a pathological measure of LMN degeneration, as previously done by Spencer *et al*^24^. Overall anterior horn cell loss was graded as mild, moderate, marked, or severe. Marked anterior horn cell loss was the most represented finding across both ALS types (58.3% in ALS-TDP-43(-), 75% in ALS-TDP-43(+)). Although the distribution of ALS-TDP-43(+) cases indicated a greater severity of cell loss in the anterior horn compared to ALS-TDP-43(-) cases, the association was not statistically significant (Wilcoxon rank-sum test, p = 0.081, **Fig. 3D**). An ordinal logistic regression was run to determine whether ALS-TDP-43(+) cases were likely to show more severe cell loss in the anterior horn than ALS-TDP-43(-) cases. Although a higher odds ratio was estimated (OR = 5.21), this association was trending, but not statistically significant (p = 0.083).

Postmortem tissue quality was assessed at the VA Boston Health care System prior to our involvement (**Table 4)**. The reported postmortem interval (PMI) metrics did not differ significantly between groups (PMI-cr: p= 0.271; PMI-frz: p= 0.076). Frozen occipital tissue from each case was utilized to assess pH, which can provide insight into agonal state^85^, as well as RNA integrity number (RIN)^86,87^ and DV200^88^, tissue quality measures that tend to decrease over time after death. We found no significant differences in occipital pH (p = 0.503), DV200 (p = 0.441), or RIN (p = 0.149), indicating comparable tissue quality and state between groups.

**Table 4.** Quality measures of tissue from study participants.

| Characteristic | Overall | Control | ALS,<br>TDP-43(-) | ALS,<br>TDP-43(+) |
| --- | --- | --- | --- | --- |
|  | *n* = 36 | *n* = 12 | *n* = 12 | *n* = 12 |
| **PMI-cr (time to body on ice), hours** |  |  |  |  |
| Mean (± SD) | 3.8 ± 3.9 | 2.7 ± 1.4 | 4.4 ± 3 | 4 ± 6.2 |
| Median (IQR) | 2 (1.2, 5.8) | 2 (1.9, 3.2) | 4.75 (1.5, 6.5) | 0.965 (0.3, 4.8) |
| Range | 0–14.25 | 1.5–5 | 1–9.25 | 0–14.25 |
| n included | 27 | 8 | 11 | 8 |
| **PMI-frz (time to brain frozen), hours** |  |  |  |  |
| Mean (± SD) | 45.1 ± 26.9 | 65.1 ± 42.3 | 40.5 ± 14.5 | 34.6 ± 10.7 |
| Median (IQR) | 37.7 (28.9, 48) | 50 (39.5, 99.3) | 37.5 (29.7, 47.1) | 33.9 (27.9, 37.9) |
| Range | 3–128 | 3–128 | 25–73 | 22.58–63.75 |
| n included | 33 | 9 | 12 | 12 |
| **pH, occipital** |  |  |  |  |
| Mean (± SD) | 6.3 ± 0.4 | 6.3 ± 0.6 | 6.2 ± 0.4 | 6.4 ± 0.4 |
| Median (Q1, Q3) | 6.3 (6.0, 6.6) | 6.3 (6.0, 6.6) | 6.2 (6.0, 6.6) | 6.5 (6.2, 6.7) |
| Range | 5.2–7.4 | 5.2–7.4 | 5.6–6.9 | 5.8–6.9 |
| n included | 32 | 10 | 12 | 10 |
| **RNA Integrity Number (RIN), occipital** |  |  |  |  |
| Mean (± SD) | 5.9 ± 1.3 | 5.6 ± 1.3 | 5.4 ± 1.5 | 6.5 ± 0.9 |
| Median (Q1, Q3) | 6.2 (5.0, 6.7) | 5.85 (5.0, 6.5) | 5.8 (4.6, 6.6) | 6.6 (6.2, 6.9) |
| Range | 2.8–8 | 2.8–7.1 | 2.8–7.1 | 4.8–8 |
| n included | 34 | 10 | 12 | 12 |
| **RIN, PosF** |  |  |  |  |
| Mean (± SD) | 5.8 ± 1.6 | 6 ± 1.5 | 5.6 ± 1.5 | 5.9 ± 2 |
| Median (Q1, Q3) | 6.4 (5.4, 7.1) | 6.5 (5.8, 7.2) | 6.2 (4.5, 6.5) | 6.45 (5.55, 7.5) |
| Range | 2.1–7.9 | 3–7.3 | 2.5–7.1 | 2.1–7.9 |
| n included | 33 | 11 | 12 | 10 |
| **RIN, SC-C** |  |  |  |  |
| Mean (± SD) | 5 ± 2.2 | 4.1 ± 2.7 | 5.3 ± 1.4 | 5.7 ± 2.1 |
| Median (Q1, Q3) | 5.55 (4.6, 6.6) | 5.4 (2.1, 5.8) | 5.2 (4.6, 6.0) | 6.4 (5.2, 7.2) |
| Range | 0–7.8 | 0–6.8 | 2.7–7.8 | 0–7.4 |
| n included | 34 | 11 | 12 | 11 |
| **RIN, SC-L** |  |  |  |  |
| Mean (± SD) | 4.9 ± 1.5 | 5.1 ± 1.6 | 4.9 ± 1.2 | 4.9 ± 1.8 |
| Median (Q1, Q3) | 5.1 (4.2, 6.0) | 5.45 (4.7, 6.1) | 5 (4.0, 5.6) | 5.35 (4.2, 5.9) |
| Range | 1.4–7.2 | 1.6–6.8 | 3.1–7.2 | 1.4–7.2 |
| n included | 34 | 10 | 12 | 12 |
| **DV200, occipital** |  |  |  |  |
| Mean (± SD) | 83.9 ± 12 | 83 ± 18.7 | 83.6 ± 9.5 | 84.9 ± 6.7 |
| Median (Q1, Q3) | 88 (79, 92) | 92 (79, 95) | 87 (79.8, 89) | 87 (80, 89.8) |
| Range | 45–97 | 45–97 | 66–94 | 72–93 |
| n included | 29 | 9 | 12 | 10 |
| **DV200, PosF** |  |  |  |  |
| Mean (± SD) | 80.4 ± 7.5 | 82.3 ± 6 | 81.8 ± 6.3 | 77.3 ± 9.2 |
| Median (Q1, Q3) | 82.9 (76.6, 85.3) | 83.6 (81.0, 85.3) | 82.4 (78.1, 86.3) | 77.9 (71.5, 84.7) |
| Range | 61.1–90.19 | 69.44–90.18 | 68.29–90.19 | 61.16–88.6 |
| n included | 35 | 11 | 12 | 12 |
| **DV200, SC-C** |  |  |  |  |
| Mean (± SD) | 79 ± 10.4 | 74 ± 10.1 | 81.8 ± 5.8 | 81.1 ± 13.4 |
| Median (Q1, Q3) | 80.9 (77.4, 87.1) | 78.2 (67.1, 81.0) | 80.9 (78.6, 86.3) | 88 (80.8, 88.8) |
| Range | 51.71–92.15 | 52.13–83.6 | 71.83–92.15 | 51.71–90.25 |
| n included | 34 | 11 | 12 | 11 |

**DV200, SC-L**

|  |  |  |  |  |
| --- | --- | --- | --- | --- |
| Mean (± SD) | 77.4 ± 6.2 | 74.3 ± 6.3 | 77.8 ± 3.8 | 79.6 ± 7.2 |
| Median (Q1, Q3) | 77.3 (73.5, 81.5) | 74.1 (71.4, 78.3) | 77.3 (75.6, 80.1) | 80.7 (74.2, 84.7) |
| Range | 63.66–91.24 | 63.66–84.33 | 71.78–84.68 | 67.9–91.24 |
| n included | 34 | 10 | 12 | 12 |

**Abbreviations:** PMI, postmortem interval; RIN, RNA integrity number; DV200, percentage of RNA fragments greater than 200 nucleotides. **Note:** PMI was defined as the time from death to either placement of the brain on ice or frozen. Summary statistics were calculated using available data. Missing values were excluded from calculations.

Prior to transcriptomic analyses, we assessed region-specific differences in RIN and DV200 analyses and compared these measures to those established by the VA using occipital tissue. RIN demonstrated a significant effect of region (aligned rank transform [ART] ANOVA, p = 0.00003, **Fig. 3E**), but not disease group (p = 0.477) or the GroupxRegion interaction (p = 0.237). Holm-adjusted posthoc assessments revealed significantly lower RIN in SC-C (p = 0.04) and SC-L (p = 0.001) tissues compared with occipital tissue RIN, however the difference between posterior frontal cortex (PosF) and occipital RIN was not significant (p = 0.499). PosF also demonstrated significantly higher RIN than SC-C (p = 0.008) and SC-L (p = 0.0001). No significant difference was observed between SC-C and SC-L (p = 0.3730)

Analyses of DV200 variance similarly revealed a significant effect of region (p = 0.0004, **Fig. 3F**), but not disease group (p = 0.699). However, the GroupxRegion interaction was significant (p = 0.03), indicating regional differences in DV200 varied by diagnosis. Holm-adjusted posthoc tests revealed significantly lower DV200 in SC-L compared to occipital tissue (p = 0.0001), however trending decreases were observed in PosF (p = 0.0584) and SC-C (p = 0.0584). No other significant differences were detected between regions. Because of the significant Group x Region interaction, regional differences were also examined within each diagnostic group; however, no individual comparison remained significant following Holm correction.

Although tissue quality measures varied between regions, a significant effect between disease groups was not detected. Overall, RNA quality was determined to be comparable between diagnostic groups, supporting the reliability of transcriptomic analyses within a region.

### TDP-43 status shows region-specific neuroinflammation, synaptic, and metabolic changes in ALS

Targeted transcriptomic profiling was performed on PosF, SC-C, and SC-L tissues, using the Human PanCancer IO 360 CodeSet. To further assess transcriptomic differences between these tissues, we utilized the four specimens with the highest DV200 values for each region for additional neuropathology-focused profiling via the Human Neuropathology CodeSet. All samples passed our predefined QC criteria (see **Methods**), and CodeSet normalization factors fell within our acceptable 0.1-10 range. The Human PanCancer IO 360 CodeSet displayed a range of ∼0.34-8.6, while the Human Neuropathology Codeset ranged from ∼0.5-1.7. Genes detected in ≥20% of samples within at least one disease condition group were retained for downstream analyses.

We first asked whether transcriptional differences could be detected across disease groups within posterior frontal cortex (PosF) and spinal cord (cervical and lumbar levels combined) tissue separately using the Neuropath and CancerIO nCounter panels. Differential expression (DE) analysis showed that transcriptional changes differed mainly by anatomical region and TDP-43 status, suggesting that ALS is associated with distinct molecular states rather than a uniform disease-associated response. The most prominent TDP-43-associated signatures were observed in PosF within the Neuropathology panel, where 29 genes were differentially expressed between ALS-TDP-43(+) and control cases (**Fig. 4a**). The expression pattern in ALS-TDP-43(+) cases showed upregulation of immune and stress-associated genes compared to control cases, including *ITGAX, TREM2, CD68, APOE*, and *HMOX1*, accompanied by downregulation of genes involved in neuronal and synaptic function. Significant pathways enriched in the KEGG and Reactome databases confirmed the DE results and showed enrichment of cytokine and cell-surface interaction programs in ALS-TDP-43(+) samples, whereas pathways associated with the neuronal system, synaptic transmission, and glutamatergic and NMDA receptor signaling were negatively enriched (**Fig. 4b)**. Together, these findings indicate that TDP-43 pathology in PosF is associated with a shift toward an immune-reactive tissue environment and loss of neuronal and synaptic pathways. No genes were significantly differentially expressed between ALS-TDP-43(-) and control in PosF samples on either panel, indicating that the prominent cortical transcriptional changes were mostly associated with the ALS-TDP-43(+) group. (**Supplementary File 1**).

**Figure 4.**
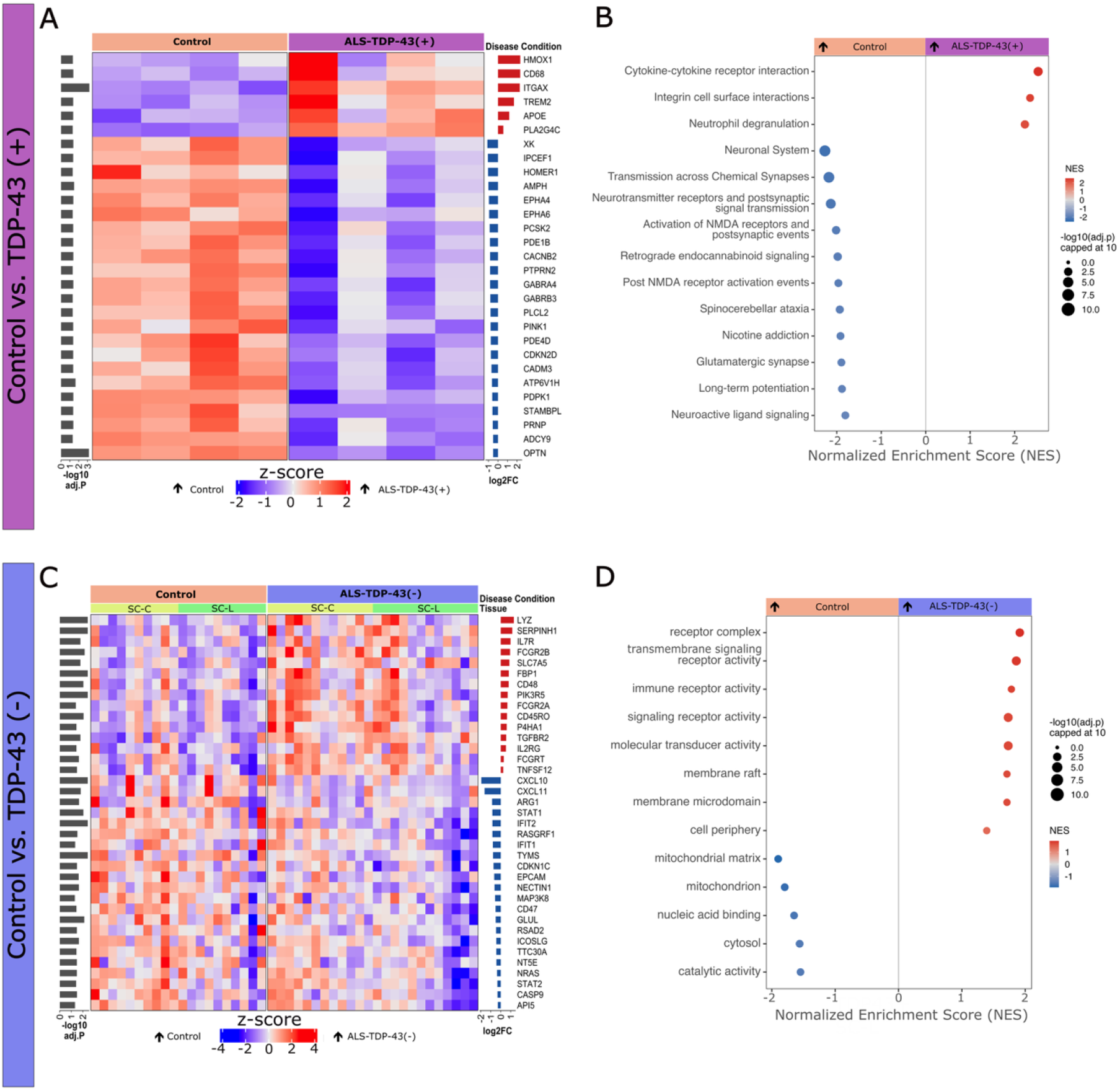
TDP-43 pathology showed region-specific molecular signatures: synaptic loss with immune activation in motor cortex, and receptor-signaling induction with mitochondrial suppression in spinal cord. **(A)** Heatmap of significantly differentially expressed genes (DESeq2 multi-factor model, Benjamini-Hochberg–adjusted p < 0.05) for ALS-TDP-43(+) vs. Control for Neuropath panel in posterior frontal cortex (PosF) samples. **(B)** Gene set enrichment analysis (GSEA) bubble plot summarizing enriched terms from KEGG and Reactome for the ALS-TDP-43(+) vs. Control comparison in PoSF samples. **(C)** Heatmap of significantly differentially expressed genes for ALS-TDP-43(-) vs. Control for CancerIO panel in spinal cord (SC-C and SC-L) samples. **(D)** GO Molecular Function and GO Cellular Component— pooled onto one shared axis per panel and ranked by normalized enrichment score (NES). For heatmaps (**A,C**), columns are annotated by Disease Condition (and additionally by Tissue in **C**); rows are genes, row-scaled to z-score and sorted into two blocks by direction of change (up, red; down, blue). Side bars show each gene’s log2 fold-change (right) and −log10(adjusted p-value) (left). For GSEA plots (**B,D**), point color indicates NES (red, enriched in the TDP43 disease group; blue, enriched in Control) and point size indicates −log10(Benjamini-Hochberg–adjusted p-value), capped at 10. Terms shown pass adjusted p < 0.05. Full GSEA results across all five databases tested (GO Biological Process, Molecular Function, Cellular Component, KEGG, Reactome) are provided in **Supplementary File 3.**

On the other hand, a different molecular phenotype was observed in the spinal cord. Within the CancerIO panel, 37 genes were differentially expressed between ALS-TDP43(-) and control samples (**Fig. 4c**). These changes reflected broad remodelling of immune and metabolic processes. Genes involved in immune-receptor and cell-surface signaling were increased in ALS-TDP-43(-) cases compared to controls, whereas several interferon-associated transcripts, including *CXCL10*, *CXCL11*, *STAT1*, and *IFIT2*, were reduced. Significant pathways identified in the GO:MP and GO:CC databases similarly showed positive enrichment of receptor-complex, immune-receptor, and cell-periphery functions, as well as negative enrichment of mitochondrial and catalytic processes (**Fig. 4d**). Thus, ALS spinal cord lacking detectable TDP-43 pathology exhibited substantial molecular remodelling involving immune signaling and cellular metabolism. By comparison, the ALS-TDP-43(+) versus control contrast identified a much smaller CancerIO signature, while no significant genes were detected in spinal cord using the Neuropathology panel (**Supplementary File 2**).

Together, these findings suggest that ALS engages at least two region-specific molecular programs: a) an immune-driven and synapse-suppressive program in motor cortex and b) an immune-receptor and metabolic remodelling in spinal cord that is detectable in the absence of TDP-43 pathology. We next asked whether these regional and pathological programs could be resolved at single neuron, spatial resolution.

### Spatial proteomics reveals cell-cycle and senescence-associated remodeling in ALS spinal cord neurons

We examined molecular alterations directly within neurons by performing GeoMx spatial proteomic profiling of single neuron areas of interest (AOIs) from PosF, SC-C, and SC-L. Due to a small sample size and low statistical power, no individual proteins survived the multiple-testing correction. We therefore examined coordinated pathway-level changes using gene set enrichment analysis of the full ranked protein profiles. The resulting proteomic signatures were strongly region dependent (**Fig. 5a-c**). In PosF, pathways enriched in ALS-TDP-43(+) compared to controls included those dominated by translation, RNA processing, and protein metabolism, whereas mitochondrial pathways were negatively enriched (**Fig. 5a**). In contrast, neuronal protein profiles in the ALS-TDP-43(-) spinal cord tissues compared to controls indicated strong positive enrichment of mitotic cell-cycle pathways, such as DNA replication, DNA replication pre-initiation, and M phase (**Fig. 5b**). This signature was accompanied by reduced enrichment of metabolic and mitochondrial respiratory pathways. A similar, but broader, phenotype was observed in comparing pathways enriched in ALS-TDP-43(+) spinal cord to controls, where multiple cell cycle checkpoints, such as G1/S and G2/M transitions, mitotic prophase, DNA repair, and TP53-associated regulation increased (**Fig. 5c**). Importantly, senescence-associated secretory phenotype (SASP) was also positively enriched specifically in the ALS-TDP-43(+) comparison. These findings support evidence that aberrant activation of cell-cycle– and senescence-associated programs is a prominent feature of ALS spinal cord neurons, while being considerably less evident in PosF.

**Figure 5.**
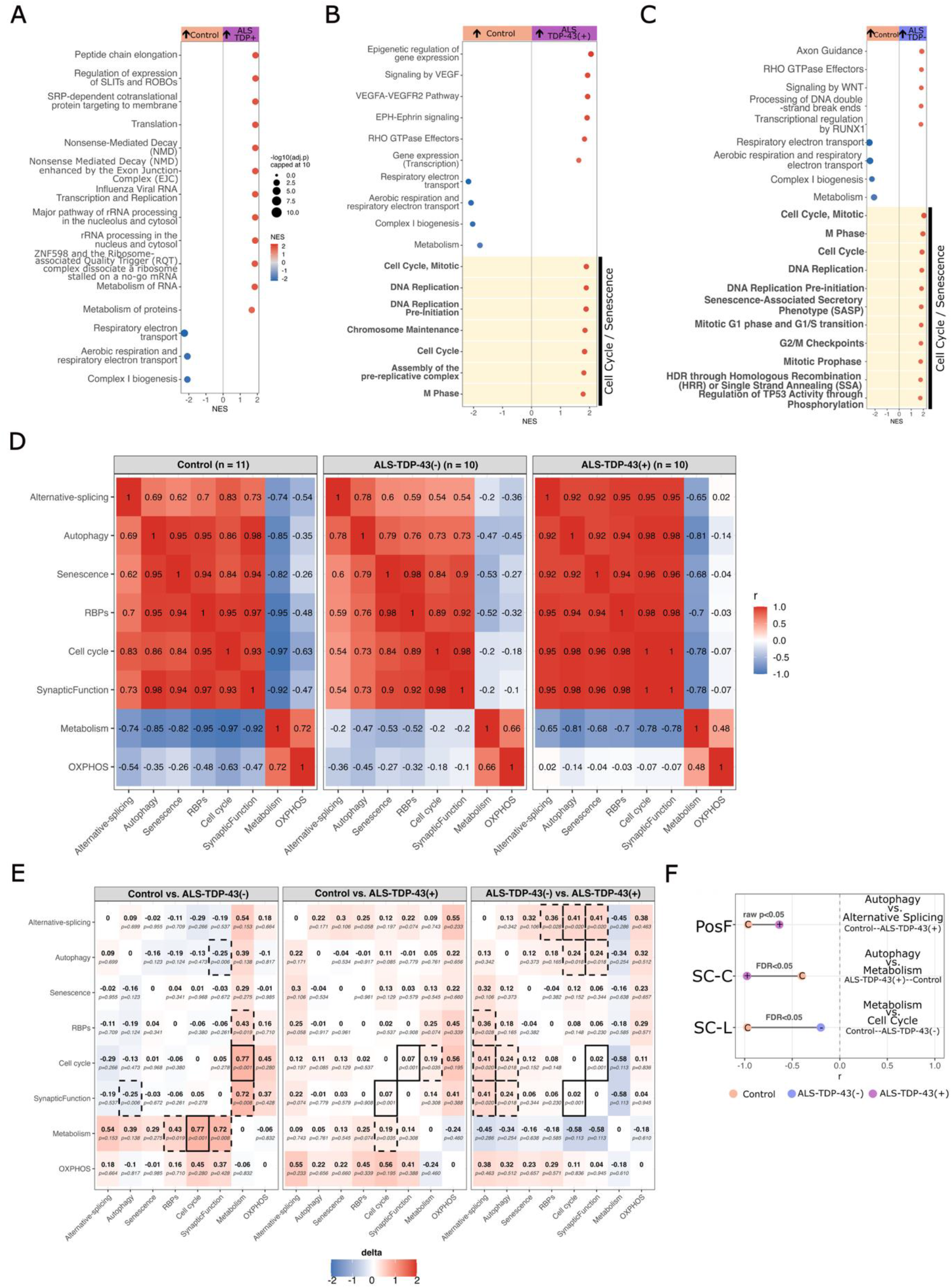
Pathway enrichment distinguishes a cell-cycle and senescence signature in spinal cord that is absent in PosF. Gene set enrichment analysis (GSEA, Reactome database) of GeoMx spatial proteomic protein rankings for three ALS-TDP-43(-) vs Control pathology contrasts. **(a)** PosF: Control vs TDP43_P. **(b)** Spinal cord (SC-C+SC-L pooled): ALS-TDP-43(-) vs Control. **(c)** Spinal cord (SC-C+SC-L pooled): ALS-TDP-43(+) vs Control. Each point is a Reactome term significant at GSEA-adjusted p < 0.05 (top 15 shown per panel); point position and color denote normalized enrichment score (NES; red/positive = up in TDP43 pathology, blue/negative = down in TDP-43 pathology relative to Control), and point size denotes −log10(adjusted p-value), capped at 10. Terms related to cell-cycle re-entry and cellular senescence are flagged with a pale-yellow background. **(d)** Full pairwise Spearman correlation matrix (r) among all eight AOI-level protein program eigenprotein scores in SC-L, computed at the case level separately within each condition (Control n=11, ALS-TDP-43(-) n=10, ALS-TDP-43(+) n=10). The color denotes the correlation coefficient itself (red = positive r, blue = negative r, diverging at 0). Two broad co-regulated blocks are visible in Control: Alternative-splicing/Autophagy/Senescence/RBPs/Cell cycle/SynapticFunction move together (r = 0.6–0.98), sharply anti-correlated with Metabolism and OXPHOS (r = −0.26 to −0.97). **(<u>e</u>)**Change in pairwise co-regulation between all eight AOI-level protein program eigenprotein scores in SC-L for three group-pair contrasts. Cell color denotes Δr direction; solid border = BH-adjusted p < 0.05; dashed border = nominal (unadjusted) p < 0.05 only. Metabolism–Cell cycle shows the largest, most significant shift of any program pair (Δr = 0.77, p < 0.001, Control vs ALS-TDP-43(-)) — already present at the ALS-TDP-43(-) stage. **(f)** The single strongest program co-regulation shift identified in each tissue (Spearman r, case-level program eigenprotein scores, baseline Control to the comparison group with strongest evidence): PosF (Autophagy vs Alternative-splicing, Control vs ALS-TDP-43(+), nominal p < 0.05), SC-C (Autophagy vs Metabolism, Control vs ALS-TDP-43(+), FDR < 0.05), and SC-L (Metabolism vs Cell cycle, Control vs ALS-TDP-43(-), FDR < 0.05, matching panel d). Point color denotes disease condition (Control/ALS-TDP-43(-)/ALS-TDP-43(+)).

The presence of cell-cycle enrichment in spinal cord from both ALS-TDP43(-) and ALS-TDP-43(+) cases compared to controls further indicated that these changes were not restricted to neurons from cases with pTDP-43 pathology. Therefore, we next asked if the observed pathways are co-activated within ALS cases. We introduced eight biologically relevant modules including senescence, cell cycle, autophagy, metabolism, oxidative phosphorylation, RNA-binding proteins, alternative splicing, and synaptic function (**Table 2**). These modules were selected to align with the main findings of GSEA. We calculated eigenproteins for each program separately and compared them across disease groups (**Fig. 5d-f**). In SC-L from controls, alternative splicing, autophagy, senescence, RNA-binding proteins (RBPs), cell-cycle, and synaptic programs were strongly positively coordinated, and they all anti-correlated with Metabolism and OXPHOS (r as low as −0.97) (**Fig. 5d**). This observation showed that under normal conditions, cell-cycle-related and metabolic and oxidative programs are essentially mutually exclusive within the same tissue. This anti-correlation reduced specifically in SC-L from ALS-TDP-43(-) cases; Metabolism and Cell cycle shifted from r = −0.97 in Controls to r = −0.20 in ALS-TDP-43(-) (Δr = 0.77, FDR-adjusted p < 0.01; **Fig. 5e**). This shift was not significantly different in SC-L from ALS-TDP-43(+) cases compared to Controls (r = −0.78, nominal p = 0.035, not significant after FDR correction). Comparably, in ALS-TDP-43(+) cases, autophagy and alternative-splicing in PosF (nominal p < 0.05) and autophagy and metabolism in SC-C were mainly decoupled (FDR-adjusted p < 0.05) (**Fig. 5f**).

Together, the protein data reinforced the region-specific model raised by the transcriptional data that spinal cord molecular dysfunction occurs independently of the histopathological hallmark used to stage TDP-43 pathology, whereas cortical vulnerability is more tightly coupled to overt pathology itself. These results also identified the spinal cord as the primary site of aberrant reactivation of cell-cycle machinery in neurons. In post-mitotic neurons, activation of cell cycle checkpoints does not generally represent productive proliferation; instead, it can occur in response to cellular stress. Cell-cycle re-entry in post-mitotic neurons in response to oxidative stress has been characterized in the context of neurodegenerative diseases including Alzheimer’s disease, where aberrant cell-cycle re-entry in aging and disease can lead to either cell death or senescence^89^.

### Phospho-TDP43 shows greater disease-dependent rewiring of neuronal stress networks than total TDP43

Whether cases were identified as being in the TDP-43 positive or negative groups depended on whether the pTDP-43 inclusions were identified at any level of the brain or spinal cord in areas affected by ALS. Accordingly, we were interested in whether the pathological marker itself — phosphorylated TDP-43 (pTDP-43, S409/S410) — and total TDP-43 protein varied by disease condition (Control/ALS-TDP-43(-)/ALS-TDP-43(+)) in each tissue (**Fig. 6a–c**). The two markers diverged in a region-specific way.

**Figure 6.**
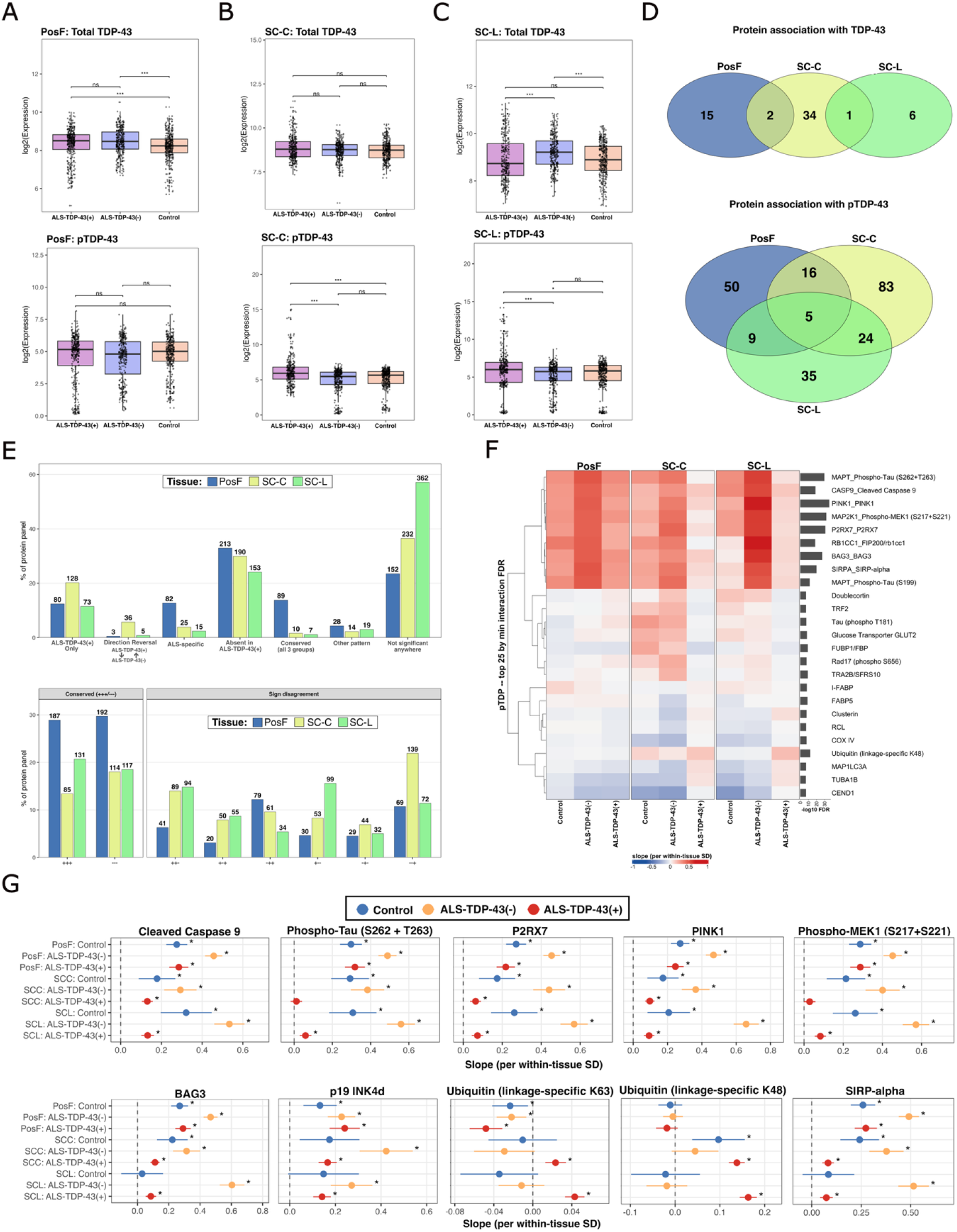
Within-case pTDP-43 and TDP-43 protein associations are disease-stage-dependent and vary by anatomical region. **(A–C)** Native TDP-43 (top) and phospho-TDP-43 (S409+S410; bottom) protein levels (log2, AOI-level) by disease condition (Control/ALS-TDP-43(-)/ALS-TDP-43(+)) in **(A)** motor cortex (PosF), **(B)** cervical spinal cord (SC-C), and **(C)** lumbar spinal cord (SC-L). Boxes show median ± IQR with individual AOIs overlaid; brackets show pairwise Wilcoxon rank-sum tests, BH-adjusted within each panel (ns, not significant; *P<0.05; **P<0.01; ***P<0.001). **(D)** Overlap across tissues of proteins whose within-case association with pTDP-43 (top) or TDP-43 (bottom) is significant only in TDP43+ neurons (“ALS-specific”), from mixed-effects models fit separately per tissue (signal×DemoGroup interaction, random effect (1|CaseNumber), AOI-loading– and AOI-area-corrected). Circles are colored by tissue; numbers are protein counts in each region. **(E)** Distribution of disease-stage archetypes (top) and raw Control/ALS-TDP-43(-)/ALS-TDP-43(+) sign patterns (bottom) across the tested protein panel, by tissue, for the pTDP-43 signal. The order of signs is PosF, SC-C, and SC-L. Archetypes summarize each protein’s pattern of significance and sign across the three groups (e.g., “ALS-TDP-43(+) Only” = significant only in TDP43+; “Conserved” = same significant sign in all three); bars show percent of that tissue’s panel, with protein counts labeled. **(F)** Top 25 proteins by minimum Control/ALS-TDP-43(-)/ALS-TDP-43(+) interaction FDR across tissues, showing each protein’s within-case pTDP-43 slope (standardized per within-tissue SD) in all nine Tissue×Disease Condition cells. Right annotation shows −log10(interaction FDR); rows are clustered by response shape (Pearson correlation across cells). **(G)** Effect size (â) ± 95% CI for a curated panel of candidate proteins across all nine Tissue×Disease Condition cells, from the same mixed-effects models as (d–f). Asterisks mark BH-adjusted P<0.05 for that cell’s own contrast.

In PosF (**Fig. 6a**), total TDP-43 was significantly elevated in both ALS-TDP43(-) and ALS-TDP-43(+) cases relative to Controls (p<0.0001) but did not differ between the two disease groups. Meanwhile, pTDP-43 itself showed no significant differences across any group. In spinal cord (**Fig. 6b-c**), the pattern was essentially reversed, with total TDP-43 being largely stable across groups and pTDP-43 significantly elevated in ALS-TDP-43(+) relative to both ALS-TDP43(-) and Control cases in SC-C (**Fig. 6b**, p<0.001) and SC-L (**Fig. 6c**, p<0.05–0.001). Therefore, the pathological (phosphorylated) form of TDP-43 track ALS-TDP-43(+) status directly in spinal cord, while in cortex, the disease-associated change was carried by total TDP-43 protein rather than its phosphorylated form.

To move beyond between-group comparisons, we next asked what happens at the level of the individual neuron. We tested whether an AOI’s own pTDP-43 or TDP-43 signal, expressed relative to that case-and-tissue’s mean, predicted its molecular phenotype. We modeled the association between protein abundance and within-case variation in pTDP-43 while simultaneously accounting for total TDP-43. This way, we distinguished molecular changes associated with the pathological phosphorylated form from those accompanying variation in total TDP-43 abundance (**Fig. 6d–g**). Proteins whose association with pTDP-43 was detected specifically in ALS-TDP-43(+) cases comprised 80 proteins in PosF (12.4% of the measured panel), 128 in SC-C (20.2%), and 73 in SC-L (11.5%) (**Fig. 6d**). Most of these associations were region specific, with only five proteins shared across all three regions. In contrast, substantially fewer proteins showed ALS-TDP-43(+)-specific associations with total TDP-43—17 in PosF, 37 in SC-C, and 7 in SC-L—indicating that the extensive disease-dependent remodeling was preferentially associated with pTDP-43 rather than total TDP-43 abundance.

The broader distribution of association patterns reinforced this distinction (**Fig. 6e**). For pTDP-43, only 13.8% of proteins in PosF and approximately 1% in either spinal cord region maintained a conserved association across Control, ALS-TDP43(-), and ALS-TDP-43(+) cases. Instead, a large fraction followed the predefined “*compensatory-lost-in-ALS-TDP-43(+)*” pattern, in which an association present in neurons from Control and/or ALS-TDP-43(-) cases was absent in ALS-TDP-43(+) neurons. This pattern accounted for 32.9% of proteins in PosF, 29.9% in SC-C, and 24.1% in SC-L. By comparison, associations with total TDP-43 levels were predominantly conserved across disease groups, accounting for 69.1%, 51.8%, and 67.8% of proteins in PosF, SC-C, and SC-L, respectively (**Fig. S2**). Thus, phosphorylation of TDP-43 was associated with substantially greater reorganization of the neuronal proteomic state than variation in total TDP-43 itself.

Proteins with the strongest disease-dependent pTDP-43 associations were relevant to neuronal stress, mitochondrial quality control, proteostasis, cell-death signaling, and neurodegeneration pathways (**Fig. 6f–g**). Phospho-tau (S262/T263), PINK1, BAG3, cleaved CASP9, P2RX7, phospho-MEK1 showed strong positive relationships with pTDP-43 in several Control and ALS-TDP-43(-) groups, specifically in spinal cord from ALS-TDP43(-) cases. These relationships were significantly reduced in ALS-TDP-43(+) SC-C and SC-L. The cell-cycle inhibitor p19/CDKN2D also increased with local pTDP-43 burden in ALS neurons across regions, linking pTDP-43 accumulation to the cell-cycle-associated phenotype identified in the previous section. In contrast, ubiquitin-associated responses became more associated with pTDP-43 in ALS-TDP-43(+) spinal cord. K48– and K63-linked ubiquitin showed strong positive associations with pTDP-43 specifically in ALS-TDP-43(+) SC-C and SC-L (**Fig. 6g**). This shift suggests that as pTDP-43 pathology becomes established, its local molecular context changes from broad coupling with mitochondrial, stress-signaling, and apoptotic pathways toward stronger engagement of ubiquitin-dependent mechanisms.

Together, these findings show proteins that covary with pTDP-43 are extensively reorganized according to anatomical region and TDP-43 pathology status. The relative preservation of total TDP-43 associations across disease groups further suggests that this remodeling is linked specifically to the pathological phosphorylated form of TDP-43. In spinal cord, the attenuation of pTDP-43 coupling with mitochondrial and stress-response proteins together with increased coupling to ubiquitin pathways is consistent with a transition in how neurons respond to accumulating TDP-43 pathology, potentially reflecting failure of adaptive stress responses and increasing dependence on proteostatic clearance mechanisms.

## Discussion

Demographic analyses reveal key differences between groups in our cohort. We identified a significantly lower age of diagnosis and longer disease duration in ALS-TDP-43(-) compared to ALS-TDP-43(+) cases. ALS-TDP-43(+) cases also showed a trend toward greater anterior horn cell loss, although this difference did not reach statistical significance. Together, these observations are consistent with distinct clinical trajectories associated with TDP-43 pathology status in this cohort. However, given the relatively small sample size and the cases reflecting the late disease stages, it is difficult to define mechanisms underlying earlier disease stages and to distinguish whether these conditions are initiated or progress via similar mechanisms.

A major challenge in understanding neurodegenerative disease is the cellular and molecular heterogeneity present in affected tissues. Bulk approaches measure molecular signals across heterogeneous populations of neurons, glia, vasculature, and other cell types, potentially diluting cell-specific signatures important to understanding mechanisms of disease. A benefit of using spatial proteomic approaches is the ability to isolate individual motor neurons to pinpoint proteomic differences in affected cells^90^, rather than pooling multiple cell-types in bulk approaches. This is particularly valuable in neurons, where nuclear profiling alone excludes much of the cytoplasmic compartment^49–51^. Our mask-based GeoMx approach captured protein signals from individual neuronal somata while minimizing contributions from surrounding neuropil and neighboring cells. This strategy is particularly relevant for identifying senescence-associated neuronal states, which may occur in a minority of neurons —estimated at only 2–3% of neurons in postmortem human brain^35^— and therefore difficult to resolve in bulk tissue.

The results of our targeted transcriptomic and proteomic profiling revealed region– and pathology-dependent phenotypes in ALS. In the spinal cord, changes in pathways associated with cell surface and cell-cycle signaling were similar between ALS cases with and without TDP-43 pathology. This suggests convergent pathway-level remodeling despite differences in TDP-43 status, indicating that TDP-43 pathological inclusions may not be required for neurons to become vulnerable to ALS-associated degeneration in the spinal cord tissue. This finding is consistent with prior literature suggesting that neuronal cell cycle dysregulation involving p16 and p21 occurs in ALS^42^. However, its presence in neurons lacking pTDP-43 pathological inclusions is particularly intriguing, given evidence linking TDP-43 loss or dysfunction to neuronal senescence^30^ and altered cell-cycle regulation^91^. TDP-43 has been shown to localize to cytoplasmic stress granules in response to cellular stress, including neuronal injury^92^. Therefore, although TDP-43 pathology is tied to a more severe disease phenotype in our cohort (as measured by disease duration and severity of anterior horn cell loss), the relationship between TDP-43 pathology and these cellular changes remains unclear. This raises the possibility that TDP-43 pathology may represent a consequence of a stressed neuronal state, rather than acting as the primary driver of these molecular changes, an idea which warrants further investigation.

ALS cases presenting with TDP-43 pathology demonstrated significant alterations to pathways associated with cellular senescence and stress responses in both the motor cortex and the spinal cord. ALS-TDP-43(+) cases presented with a reduction in pathways associated with neuronal signaling in the PosF, a feature that is intriguing given that these cases trended toward greater overall severity as measured by anterior horn degeneration. Although anterior horn degeneration does not directly reflect cortical disease, the association between pTDP-43 pathological inclusions and downregulation of pathways associated with neuronal signaling in the PosF suggest that presence of pTDP-43 could be explained by a distinct pTDP-43-associated neuronal phenotype within the motor cortex that influences downstream neuronal dysfunction. This interpretation is consistent with existing evidence that TDP-43 is a major regulator of neuronal RNAs essential for neuronal and synaptic function^91^.

Alterations in relational coupling between cellular programming in the lumbar spinal cord were identified in ALS cases, with TDP-43 status influencing the strength of correlation. ALS-TDP-43(-) cases demonstrated broad uncoupling of coordinated proteomic signaling programs that were associated in controls, with several relationships that were strongly positively or negatively correlated becoming less so. This is particularly prevalent in the association with metabolism and oxidative phosphorylation. ALS-TDP-43(+) cases, on the other hand, retained strong patterns of coordination apart from oxidative phosphorylation, which appeared to become nearly entirely uncoupled. These findings suggest that TDP-43 status may be associated with distinct molecular states as well as how cellular programs remain coordinated during disease.

The relationship between TDP-43 pathology and neuronal molecular state was supported by our comparison of total TDP-43 and pTDP-43. In the PosF, total TDP-43 was elevated in both ALS groups, whereas pTDP-43 did not significantly differ. In the spinal cord, the opposite pattern was observed; pTDP-43 was significantly elevated in the ALS-TDP-43(+) group, while total TDP-43 was consistent between groups. These regional variations in pTDP-43 levels are consistent with the presence of disease-associated pTDP-43 inclusions in ALS-TDP-43(+) cases, as expected. However, the increase in total TDP-43 in the motor cortex is intriguing, given that multiple TDP-43 RNA targets have been reported as being crucial in neural development and senescence, including *NOTCH1* and *MAPT*^93^. Overexpression of TDP-43 has been identified as negatively impacting motor function in *Drosophila* at the level of the motor neuron by increasing the number of boutons at the neuromusclular junction^94^ and in zebrafish by shortening the motor neuron^95^.

Even in the absence of pTDP-43 in the PosF of the ALS-TDP-43(-) group, we identified a significantly higher level of total TDP-43 compared to the control, which within the context of the literature, may reflect a different mechanism of TDP-43 disruption that results in axonal retraction and wasting. Future studies differentiating the role of different protein isoforms of TDP-43 would inform whether another form of TDP-43 is impacting the ALS-TDP-43(-) cases.

At the individual-neuron level, within-case pTDP-43 abundance showed substantially greater disease-group-dependent proteomic coupling than total TDP-43. Many proteins involved in mitochondrial quality control, stress signaling, proteostasis, and cell-death pathways showed associations with local pTDP-43 that were present in Controls or ALS-TDP-43(-) neurons but reduced in ALS-TDP-43(+) spinal cord. K48– and K63-linked ubiquitin showed stronger positive coupling with pTDP-43 in ALS-TDP-43(+). Together, these results indicate that the molecular context in which pTDP-43 varies differs substantially according to the pathological group. The association of p19/*CDKN2D* with local pTDP-43 further links these changes to the cell-cycle-arrest phenotype identified independently by pathway analysis.

Together, our findings support a model in which ALS motor neurons exhibit distinct proteomic changes according to the presence of pTDP-43 pathology, including alterations in neuronal stress, mitochondrial function, and cell-cycle regulation. Overall, we identified the presence of pTDP-43 as being more associated with disease-dependent molecular changes than total TDP-43, indicating pTDP-43 as a more informative marker for ALS than TDP-43 alone, among the targets tested in our panels. The convergence on cell-cycle arrest and senescence-associated pathways warrants further investigation as a potential vulnerability of surviving motor neurons, while functional studies will be required to determine whether modifying these pathways can alter neuronal dysfunction or disease progression.

## Limitations

These analyses provided important preliminary findings distinguishing possible mechanisms underlying ALS, with and without TDP-43 pathological inclusions, and highlighted significant areas for further research. However, there were several limitations associated with these data.

The predominance of male participants in this cohort reflects the demographic composition of the VABBB donation population (94.1%), but does not reflect the percentage of women in the military (17.9%, as of 2024^96^). Although ALS has been reported as appearing most commonly in non-Hispanic Caucasian men^97,98^, distinct differences in onset and diagnosis have been observed in African American individuals^99^ and women^97,100,101^. Therefore, generalizability of these findings to veterans that belong to underrepresented populations, namely women and non-Caucasians, may be limited. Understanding how ALS may progress differently in females and non-Caucasian individuals, particularly among veterans, is of crucial importance for future studies.

Another confound in our analyses is the relatively small sample size within groups, which limited our ability to meaningfully define differentially expressed proteins between groups. Although the familial cases in our ALS-TDP-43(-) cases did not present with causal *SOD1* or *FUS* mutations, the presence of familial cases in this group confound our ability to compare the groups according to TDP-43 status, since the ALS-TDP-43(+) group is entirely sporadic. The presence of familial cases in the ALS-TDP-43(-) group reflects the rarity of cases without TDP-43 pathology.

Since our experimental design utilized curated panels of genes and proteins, our analyses may not reflect a comprehensive view of differentially expressed genes, proteins, and pathways between the disease groups. In this manuscript, we utilized brain sections from tissue regions of interest that reflect end-stage ALS. Since motor neurons are characteristically lost during ALS, the remaining neurons captured at end-stage may not fully reflect molecular signals underlying vulnerability of these populations.

## Author Contributions

MEO conceived and designed the study. PHD and MSP contributed to slide selection and preparation for GeoMx data collection. TCO performed AOI selection. XS performed the transcriptomics/n counter data acquisition. SKD and PHD performed statistical analyses and data visualization. SKD, MEO, PHD, FJA, and CVL contributed to interpretation of the findings. PHD, SKD, MEO, XS, FJA, and CVL contributed to writing and revision of the manuscript.

## Acknowledgements

The authors acknowledge the individuals who generously consented to donating their neural tissue to the VA BBB, as well as their families. Human neuroscience research aimed at understanding neurological disease would not be possible without this generous contribution. We would also like to acknowledge the individuals from the VA BBB for their assistance with tissue procurement and preparation for this study.

## Data and Code Availability

Raw data and code used for analyses are available upon reasonable request to the corresponding author, Miranda E. Orr.

## Supplemental

**Table S1.** Sources, catalog numbers, and lot numbers (where available) for reagents and consumables used in preparation for this manuscript.

| **Reagent** | **Source** | **Catalog #** | **Lot #** |
| --- | --- | --- | --- |
| Citrisolv | *Decon Labs* | 1601 | 260155 |
| 200 Proof Ethanol | *Decon Labs* | 2701 | 1332617 |
| 10X Citrate Buffer | *Sigma-Aldrich* | 1003807663 | MKCX9107 |
| 10X TBS-T | *Cell Signaling Technology* | 9997S007700072027 | 77 |
| Super PAP-Pen | *Biocare Medical* | PEN1111 |  |
| SignalStain Antibody Diluent | *Cell Signaling Technology* | 8112 | 48 |
| HybriSlip Hybridization Covers | *Grace Bio-labs* | 714022 | 1024003,<br>1223067 |
| GeoMx Core Ab Mix_Hs_Ilmn | *Bruker Spatial Biology* | 121300162 | 621010 |
| GeoMx NPA Ab Mix_Hs_Ilmn | *Bruker Spatial Biology* | 121300162 | 624052 |
| GeoMx NGS Orr 21-Plex | *Bruker Spatial Biology* | custom | 624114 |
| CoraLite594-conjugated IBA1<br>Recombinant monoclonal<br>antibody | *Proteintech* | CL594-81728 | 21025494 |
| HuD Antibody (E-1) Alexa<br>Fluor 647 | *Santa Cruz Biotech* | sc-28299AF647 | A1824 |
| Alexa Fluor 647 Anti-MAP2<br>antibody | *Abcam* | ab225315 | 1023468-15 |
| CoraLite Plus 488-conjugated<br>TDP-43 Polyclonal antibody | *Proteintech* | CL488-10782 | 21024546,<br>21028464 |
| FlexQuencher for Rabbit IgG | *Proteintech* | FA591-200 | LM00000432 |
| FlexBuffer | *Proteintech* | FA090-200 | 2916948 |
| FlexLinker 2.0 for Rabbit IgG<br>CoraLite 488 | *Proteintech* | FA501-200 | LM00000432 |
| Tris Buffered Saline with<br>Tween 20 (TBS-T-10X) | *Cell Signaling Technology* | 9997S | 77, 79 |
| Paraformaldehyde 16%<br>Aqueous Solution EM Grade | *Electron Microscopy*<br>*Sciences* | 15710 |  |
| 10X PBS | *R&D Systems* | 4870-500 | P276877 |
| SYTO 83 Fluorescent Nucleic<br>Acid Stain | *Invitrogen* | S11364 | 2610305,<br>2725305 |
| GeoMx DSP Collection Plate | *Bruker Spatial Biology* | 100473 | N/A |
| GeoMx DSP Instrument Buffer<br>Kit | *Bruker Spatial Biology* | 100474 |  |
| Adhesive PCR Plate Foils | *Thermo Scientific* | AB0626 | 161821 |
| Water (for RNA Work) DEPC-treated and Nuclease-free | *Fisher Bioreagents* | BP-561-1 | 236181 |
| AeraSeal film | *Excel Scientific* | A9224-50EA | MKCT3860 |
| Twin.tec PCR Plate 96, semi-<br>skirted, colorless | *Eppendorf* | 951020303 | O220955K |

**Table S1** (cont).
| **Reagent** | **Source** | **Catalog #** | **Lot #** |
| --- | --- | --- | --- |
| GeoMx Pro Code S | *Bruker Spatial Biology* | 531-121400210 | S3250611 |
| GeoMx Pro Code T | *Bruker Spatial Biology* | 531-121400210 | T32250611 |
| GeoMx Pro Code U | *Bruker Spatial Biology* | 531-121400209 | U3250611 |
| GeoMx Pro Code V | *Bruker Spatial Biology* | 531-121400209 | V3250612 |
| GeoMx Pro Code W | *Bruker Spatial Biology* | 531-121400208 | W3250620 |
| GeoMx Pro Code Y | *Bruker Spatial Biology* | 531-121400207 | Y7240918 |
| GeoMx Pro Code Z | *Bruker Spatial Biology* | 531-121400207 | Z7240918 |
| NGS Master Mix (comes in GeoMx Pro Code Pack) | *Bruker Spatial Biology* | 531-121400207,<br>531-121400208,<br>531-121400209,<br>531-121400210 | 10279587,<br>10156545,<br>10311739 |
| 0.2mL Thin-walled 12 Tube Strips | *Thermo Scientific* | AB-1112 |  |
| Safe-Lock Tubes 1.5mL, natural | *Eppendorf* | 022363204 | K198290N |
| AMPure XP Beads for DNA Cleanup | *Beckman Coulter* | A63881 | 20961700 |
| 10mM Tris-HCL with 0.05% TWEEN-20, pH 8.0 | *TEKNOVA* | T1484 | T148519G2201 |
| Qubit assay tubes | *Invitrogen* | Q32856 | FA8222652551 |
| PhiX Control v3 | *Illumina* | 15017666 | 21026405 |
| NextSeq 1000/2000 RSB with Tween 20 | *Illumina* | 20050639 | 210067874 |
| NextSeq 1000/2000 XLEAP-SBS P2 Reagent Cartridge | *Illumina* | 20101832 | 21045059 |
| NextSeq 1000/2000 XLEAP-SBS P2 Flow Cell | *Illumina* | 20094371 | 21057179 |
| NextSeq 2000 XLEAP-SBS P4 Reagent Cartridge | *Illumina* | 20101842 | 21060894 |
| NextSeq 2000 XLEAP-SBS P4 Flow Cell | *Illumina* | 20094373 | 21079171 |
| RNeasy Midi Kit | *Qiagen* | 75144 |  |
| 2-Mercaptoethanol, ≥99.0% | *Sigma-Aldrich* | M6250 |  |
| Disposable Pellet Pestle Tissue Grinder, RNase-Free, 1.5mL | *Kimble* | 749520-0090 |  |
| PrecisionGlide 21 G x 1 in. Needle | *BD* | 305165 |  |
| nCounter Human PanCancer IO 360 CodeSet | *Bruker Spatial Biology* | XT-CSO-HI0360-12 |  |
| nCounter Human Neuropathology CodeSet | *Bruker Spatial Biology* | XT-CSO-HNROP1-12 |  |

**Table S2.** Equipment used in preparation for this manuscript.

| Item | Source | Catalog Number |
| --- | --- | --- |
| Heratherm IGS180 | *Thermo Scientific* | 51028065 |
| Assorted Easydip Slide Staining Jars | *Simport* | M900-12AS |
| Easydip Slide Staining Rack | *Simport* | M905-12DGY |
| Remote Dispenser for Ultrapure Water | *Rephile* | RSP0U000N |
| TintoRetriever Pressure Cooker | *BioSB* | BSB 7008 |
| Staintray 20 Slides Staining System | *Simport* | M920-2 |
| GeoMx Digital Spatial Profiler | *Bruker Spatial Biology* | GMX-DSP |
| VTS Variable Temperature Sealer | *Vitl Life Science Solutions* | V902001 |
| Centrifuge 5910 Ri | *Eppendorf* | 5943000131 |
| LSE mini microcentrifuge | *Millipore sigma* | CLS6770 |
| 1500 Series Class II Type A2 Biological Safety Cabinet | *Thermo Scientific* | 1523A2 |
| Mastercycler X40 | *Eppendorf* | 6381000018 |
| Mastercycler Nexus Thermal Cycler | *Eppendorf* | 6339000024 |
| VeritiPro Thermal Cycler, 96 well | *ThermoFisher Scientific* | A48141 |
| DynaMag-2 Magnet | *Invitrogen* | 12-321-D |
| Qubit 4 fluorometer, with WiFi | *Invitrogen* | Q33238 |
| NextSeq 2000 Sequencing System | *Illumina* | 20038897 |
| NanoDrop 8000 spectrophotometer | *Thermo Scientific* | ND-8000-GL |
| TapeStation 4200 | *Agilent* | G2001BA |
| nCounter SPRINT Profiler | *Bruker Spatial Biology* | 52-851418-002 |

**Table S3.** Custom protein probes for GeoMx analysis.

| Antibody target | Clone |
| --- | --- |
| BAG3 | EPR3515 |
| cGAS | PolyNBP316666 |
| FIP200 | EPR26076-87 |
| MAP6 (STOP) | STOP |
| P2RX7 | CL11508 |
| PARK2 | PolyDJ9 |
| phospho TDP-43 (Ser409/Ser410) | 1D3 |
| phospho-RIPK1 (Ser161) | 1B2G1 |
| PINK1 | DU46-1.1 |
| PLCG1 | D9H10 |
| phospho-MEK1/2 (Ser217/221) | 41G9 |
| phospho-PRAS40 (Thr246) | D4D2 |
| phospho-RIP3 (Ser164/Tyr165) | EPR23660-20 |
| RIPK3/RIP3 | OTI1B3 |
| SIRP alpha | BLR049F |
| phospho-Tau (Ser199) | EPR2401Y |
| phospho-Tau (Ser262) | EPR2454 |
| TFEB | D2O7D |
| TREM2 | HL1738 |
| CASP9 | PolyCASP9t |
| p19 INK4d (CDKN2D) | pAb_ab262871 |

**Table S4.**
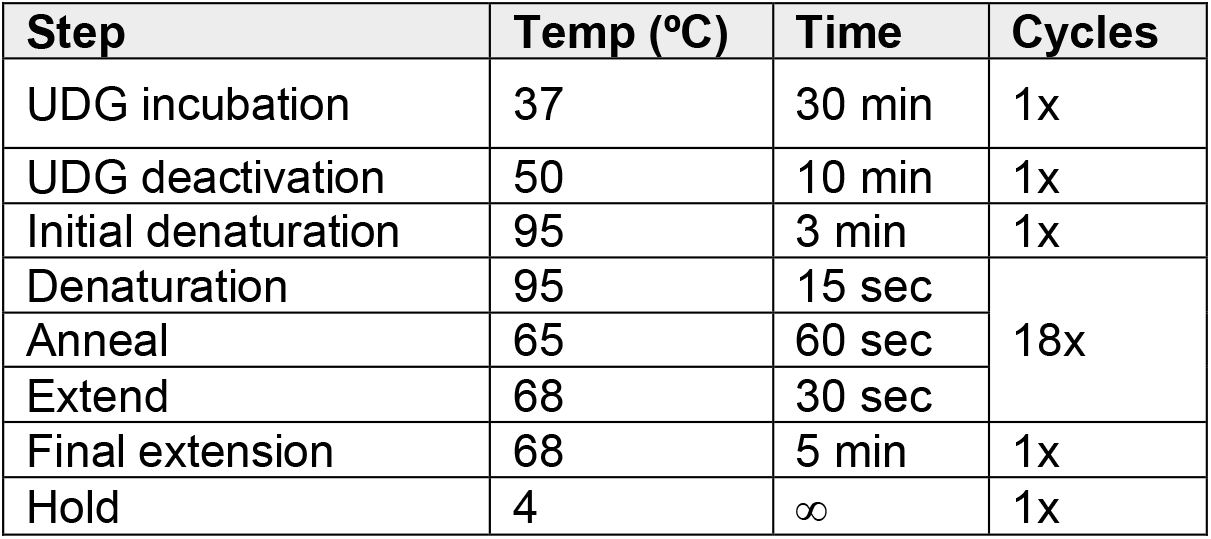
PCR setup for GeoMx DSP next generation sequencing library preparation.

| Step | Temp (°C) | Time | Cycles |
| --- | --- | --- | --- |
| UDG incubation | 37 | 30 min | 1x |
| UDG deactivation | 50 | 10 min | 1x |
| Initial denaturation | 95 | 3 min | 1x |
| Denaturation | 95 | 15 sec | 18x |
| Anneal | 65 | 60 sec |  |
| Extend | 68 | 30 sec |  |
| Final extension | 68 | 5 min | 1x |
| Hold | 4 | ∞ | 1x |

## Supplemental Figs

**Figure S1.**
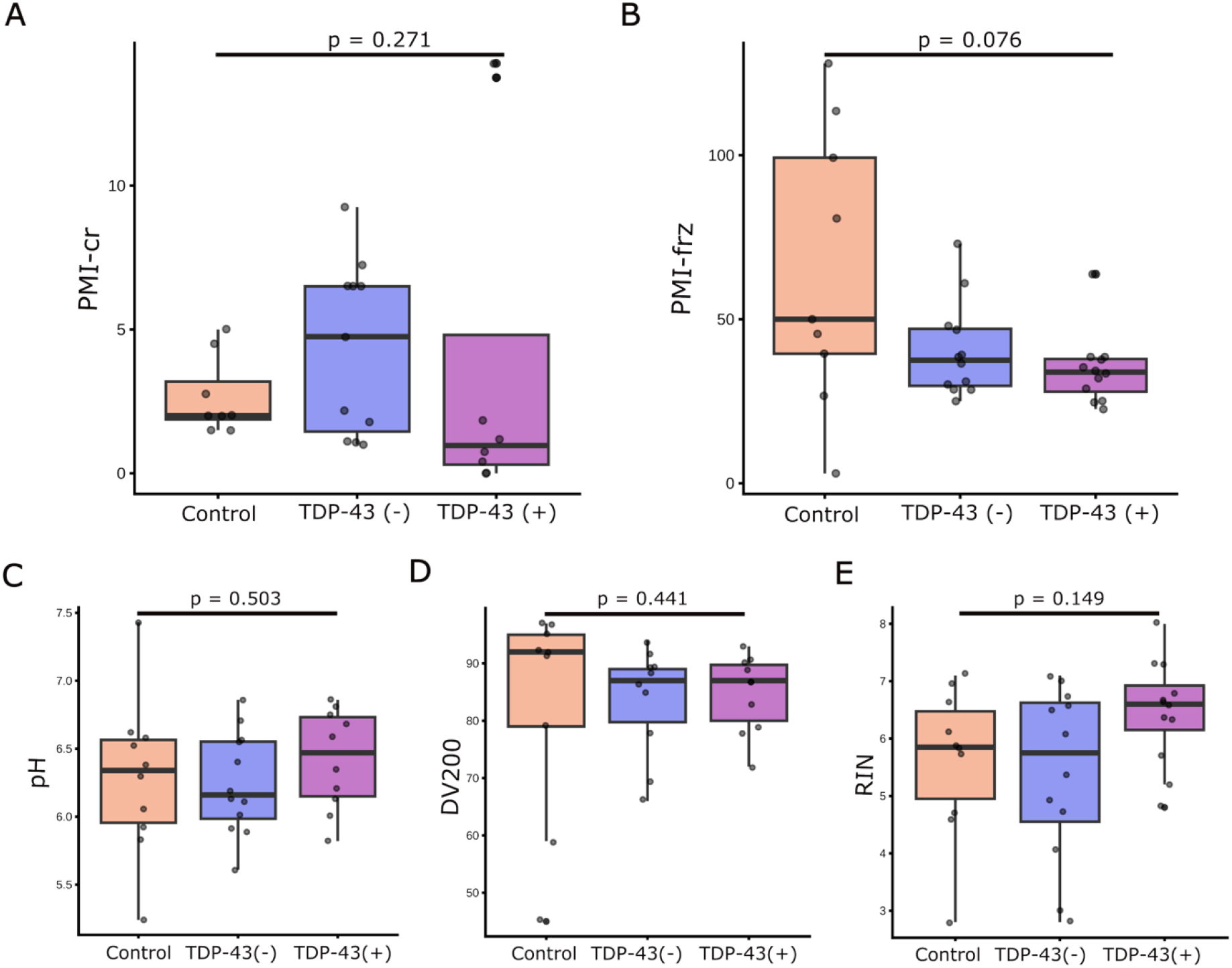
Tissue quality comparison between groups in study cohort reflect no significant differences. Box plots showing (**A**) Post mortem interval (PMI) to body on ice (PMI-cr), (**B**) PMI to brain freezing (PMI-frz) across disease status, (**C**) occipital lobe pH, (**D**) DV200, a measure representing the percentage of RNA fragments >200 nucleotides, (**E**) RNA Integrity Number (RIN) across Control, TDP43-, and TDP43+ groups. No significant differences were observed between groups for PMI-cr (Kruskal-Wallis, p= 0.271), PMI-frz (p = 0.076), pH (p = 0.503), DV200 (p = 0.441), or RIN (p = 0.149).

**Figure S2.**
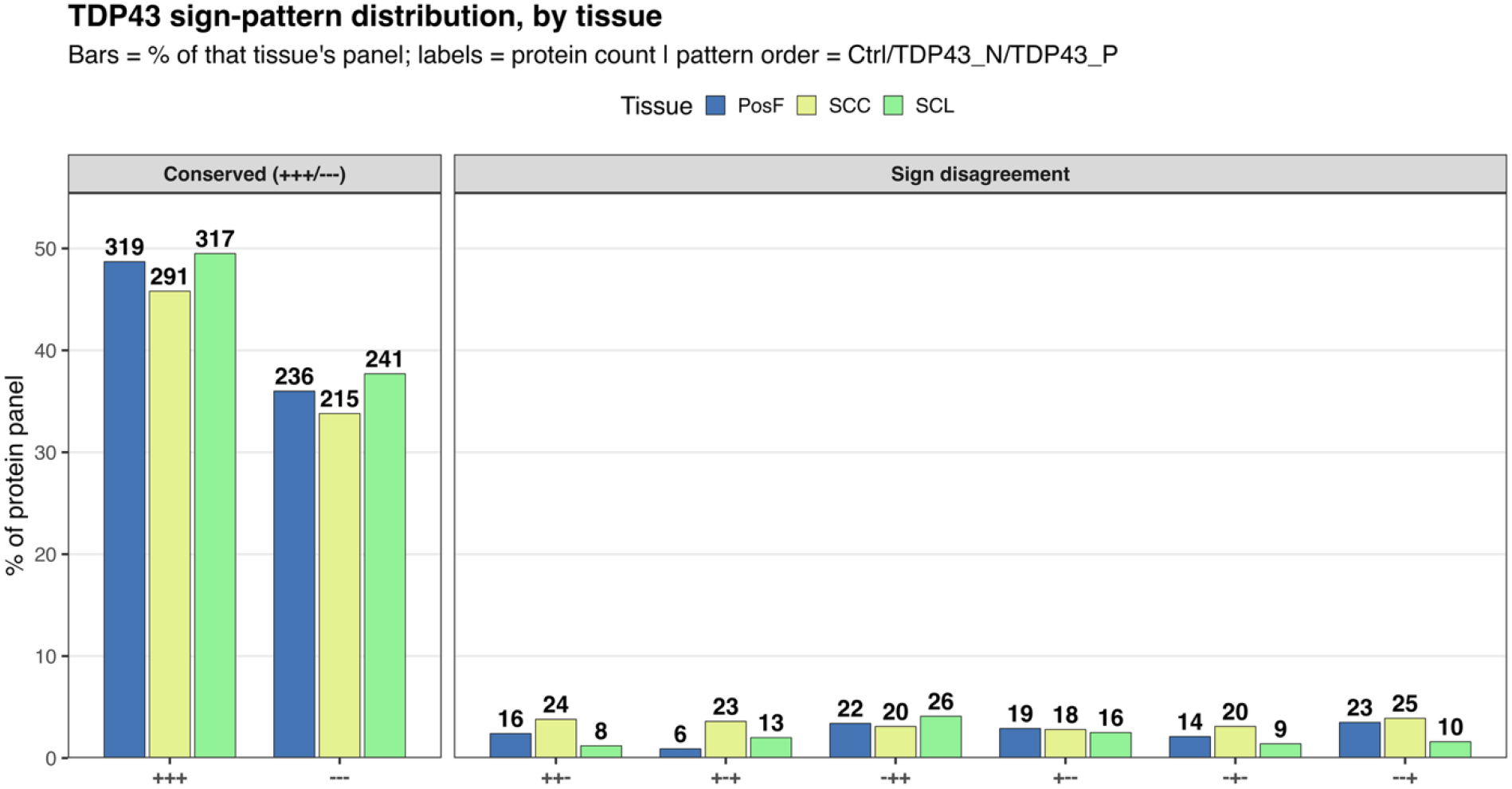
Distribution of raw Control/ALS-TDP-43(-)/ALS-TDP-43(+) sign patterns across the tested protein panel, by tissue, for total TDP-43 signal. The order of signs is as follows: PosF, SC-C, and SC-L. Bars reflect percentage of proteomic panel, with protein counts labeled.

